# Thermodynamic Stability Modulates Chaperone-Mediated Disaggregation of α-Synuclein Fibrils

**DOI:** 10.1101/2024.12.19.629136

**Authors:** Celia Fricke, Antonin Kunka, Rasmus K. Norrild, Shuang-Yan Wang, Thi Lieu Dang, Anne S. Wentink, Bernd Bukau, Alexander K. Buell

**Affiliations:** Department of Biotechnology and Biomedicine, Technical University of Denmark, Søltofts Plads, Building 227, 2800 Kgs. Lyngby, Denmark; Leiden Institute of Chemistry Leiden University Einsteinweg 55, 2333 CC Leiden, Netherlands; Center for Molecular Biology of Heidelberg University (ZMBH) DKFZ-ZMBH Alliance Heidelberg, Germany

**Keywords:** polymorphism, amyloid stability, chaperones, synuclein, Parkinson’s disease

## Abstract

The aggregation of the intrinsically disordered protein alpha-synuclein into amyloid fibrils and their subsequent intracellular accumulation are characteristic features of several neurodegenerative disorders, such as Parkinson’s disease. Currently, there are no curative treatment options available. In this study, we demonstrate that the thermodynamic stability of alpha-synuclein fibrils is a crucial factor influencing the efficiency with which they are disaggregated by the human chaperone system comprising HSP70, DNAJB1, and Apg2. We quantify the increasing stability of alpha-synuclein fibrils formed under four different solution conditions over a three-month incubation period. The chaperone system effectively disaggregates three out of the four fibril types, with varying efficiencies that correlate with their thermodynamic stability. The fibrils exhibit differential sensitivities to chaperone-mediated depolymerization, suggesting that both structural features and thermodynamic stability contribute to the susceptibility of alpha-synuclein fibrils to chaperone disaggregation. Our findings thus reveal a connection between the thermodynamic stability of fibrils and their susceptibility to chaperone-mediated disaggregation.

## Introduction

One of the hallmark characteristics of neurodegenerative diseases such as Alzheimer’s (AD) and Parkinson’s (PD) is the aberrant accumulation of protein aggregates in the form of amyloid fibrils within the brain.^[1–4]^ Although the direct causal relationship between protein aggregation and disease progression remains disputed, ^[5–7]^ the neurotoxic effects of amyloid fibrils, e.g. synaptic dysfunction and neuronal death, have been clearly demonstrated across various disease models. ^[8–11]^ Interestingly, the main protein constituents of the extracellular and intracellular deposits in Alzheimer’s disease (amyloid-beta peptide (Aβ), Tau) and Parkinson’s disease (α-synuclein (αSyn)) are also implicated in multiple other neurodegenerative disorders, each with distinct molecular and pathological signatures. ^[10,12]^ A compelling hypothesis suggests that the distinct structures of amyloid fibrils, such as those of Tau isolated from patients’ brains, could serve as signatures linked to the underlying pathology through amyloid polymorphism. ^[12,13]^ The term polymorphism refers to the ability of a single protein sequence to adopt various amyloid conformations, which differ in (i) the specific interactions within the filament core, (ii) the packing arrangement of the filament core, or (iii) the interface and organization of the protofilaments within the mature fibril.^[14–16]^

Amyloid polymorphism has been observed in most amyloid-forming proteins, including Aβ, Tau, αSyn, insulin, or glucagon.^[17– 22]^ This variability is considered a manifestation of the rugged amyloid energy landscape allowing multiple conformations to emerge and coexist under identical solution conditions, as seen for prions^[23]^, αSyn^[24,25]^, insulin^[26]^, and other amyloid proteins. Polymorphism is highly sensitive to both intrinsic factors (e.g., mutations^[27,28]^, posttranslational modifications^[29–31]^) and extrinsic factors (e.g. pH^[32]^, ionic strength^[33]^). Time is also critical, as amyloid fibrils mature structurally over months (and potentially years).^[34]^ Despite age being the highest risk factor in most of the neurodegenerative diseases^[35]^, most mechanistic aggregation studies focus on short timescales (i.e., minutes to days). Considerably less is known regarding the long-term (month to years) structural or morphological changes of amyloid fibrils, such as those observed from proteins like glucagon^[20]^, amyloid beta^[36,37]^, or αSyn^[34,38,39]^. Similarly, recent cryo-EM studies showed time-dependent development of Tau^[40]^ and IAPP^[41]^ polymorphism during their formation and maturation, clearly demonstrating that amyloid fibril evolution on timescales beyond those of the “classical aggregation assays” (typically a few days) should be considered. It is hypothesized that the “late stage” polymorphs persist in the brain due to their high thermodynamic stability. The stability varies significantly between fibrils formed by different amyloid peptides^[42]^ and between polymorphs^[43]^, with reported values ranging from -65 kJ/mol to -14 kJ/mol.^[42]^ However, only very few studies focus on the quantification of fibril stability over time.^[41]^

The structural variability of amyloid fibrils is modulated by the cellular environment and may be linked to disease pathology either directly, through structure-toxicity relationship, or indirectly, via distinct interactions with other cellular components. Molecular chaperones are essential in this context, serving as protein quality control and maintaining cellular homeostasis.^[44– 46]^ Fundamental aspects of chaperone action include assisting protein folding, preventing protein aggregation and targeting proteins to degradation systems such as the ubiquitin/proteasome system or the autophagy system.^[47–50]^ A specific chaperone system consisting of Heat Shock Protein 70 (HSP70) and co-chaperones from the J-domain protein (JDP) and Nucleotide Exchange Factor (NEF) families can disaggregate amyloid fibrils formed by αSyn^[51–54]^ Tau^[55]^, and exon1 of the Huntingtin protein.^[56]^ The mechanism is ATP-dependent and involves generating an entropic pulling force.^[51,57–59]^ HSP70 and JDP DNAJB1 mainly interact with the flexible regions in the N- and C-terminus of αSyn fibrils and not with the fibril core.^[51]^ Distinct polymorphs of αSyn fibrils disaggregate to varying degrees, which has been attributed to an interplay between the chaperones’ affinities for the fibrils and variations in the polymorph’s stabilities.^[60]^ In contrast to the chaperone binding, the stabilities have only been inferred indirectly from the fibrils’ resistance to cold denaturation, and their precise quantification is still missing. Furthermore, how amyloid thermodynamic stability, fibril polymorphism, and their temporal changes influence the efficiency of disaggregation by chaperones remains unclear.

Here, we hypothesize that the thermodynamic stability of fibrils changes as they mature with age, which in turn decreases their susceptibility towards disaggregation by the tri-chaperone system. To address this, we follow the maturation of αSyn fibrils in four different sets of solution conditions over a period of three months using a series of biochemical and biophysical assays. We observe significant condition-dependent structural and morphological changes accompanied by increasing thermodynamic stability of fibrils during their aging for some of the conditions. Furthermore, we demonstrate that the increasing stability is negatively correlated with the disaggregation efficiency of the tri-chaperone system HSP70, DNAJB1, and Apg2, for three out of four cases studied here. The correlations are sensitive to the specific fibril types, indicating that apart from the thermodynamic stability of the fibrils, their structural or morphological features influence the action of the chaperones. Altogether, our results provide an important link between fibril polymorphism, stability, and chaperone disaggregation efficiency.

## Results

### Selection of assembly conditions and experimental design

To generate distinct αSyn polymorphs, we selected four experimental solution conditions that have been used previously to obtain well-defined sets of fibril morphologies *in vitro* (Table 2).^[33,61–63]^ Specifically, these involve physiological pH and ionic strength (50 mM Tris-HCl 150 mM KCl, pH 7.4), low ionic strength (5 mM Tris-HCl pH 7.4), mildly acidic pH (20 mM MES, 150 mM NaCl, pH 6.5), and mildly basic pH (20 mM KPO4, pH 9.1) that yield assemblies termed Fibrils (Fm), Ribbons (Ri), Fibrils-65 (F65) and Fibrils-91 (F91), respectively.^[62]^ Two polymorphic structures, 2a (pdb id 6ssx) and 2b (pdb id 6sst), have been resolved in the Fm samples using cryo-EM^[64]^, whereas only a protein secondary structure assignment is available for Ribbons^[63,65]^ and F91^[66]^ based on solid-state NMR experiments. The structure of F65 is unknown but their twisted morphology, and proteinase K digestion fingerprint resemble those of the Fm^[33,62]^. Despite detailed characterization of the polymorphs, the protocols describing their assembly differ between individual publications and (based on personal communication) among different laboratories in parameters including αSyn purification, incubation time, initial monomer concentration, and absence or presence of beads during sample agitation.^[33,61–63]^ Although seemingly minor details, these parameters may significantly influence the time-dependent polymorphic distribution of the fibril preparations and, consequently, results of the downstream assays. Here, we prepared the fibrils in the four conditions described above at the same initial protein concentration, temperature, shaking speed, and sample volumes to minimize potential variation stemming from these sources (Figure 1). We characterized the time-dependent changes of αSyn polymorphism at six different time-points between 3 days and 3 months in the four experimental conditions using a series of biophysical assays (Figure 1). Next, we used chemical depolymerization and transient incomplete separation to determine the thermodynamic stability of the fibrils.^[43]^ Finally, we performed chaperone disaggregation assays using HSP70, DNAJB1, and Apg2 to investigate the relation between fibril stability and propensity for disaggregation by chaperones.

**Table 1.**
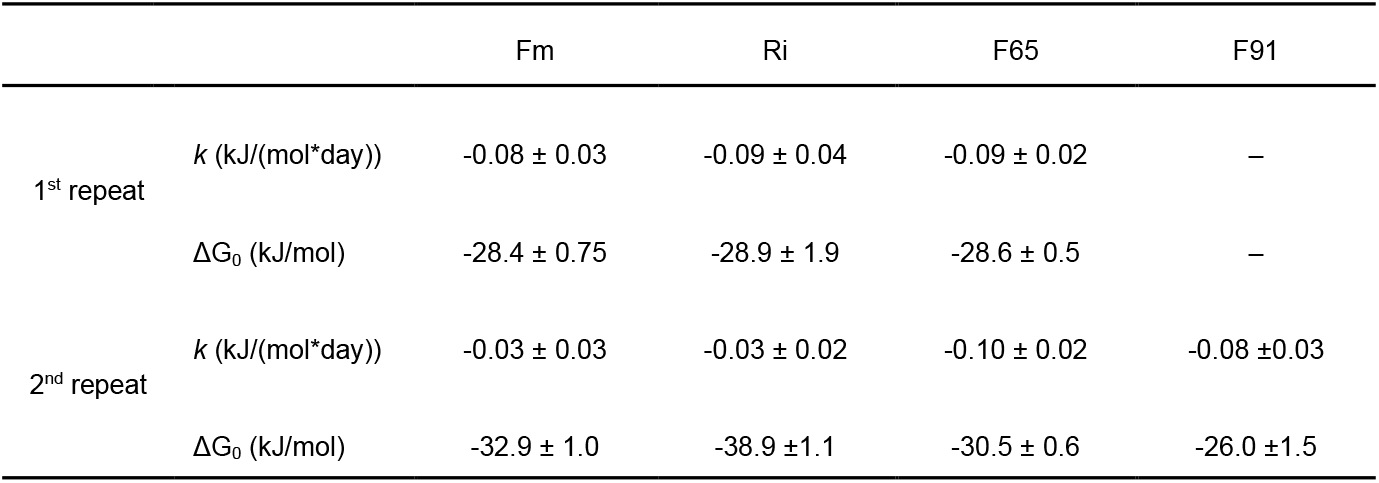
Parameters obtained from global fitting of the chemical depolymerization datasets into the linear maturation model. k – slope (rate), ΔG_0_ – intercept. m-values can be found in Supplementary Table 3.

|  |  | Fm | Ri | F65 | F91 |
| --- | --- | --- | --- | --- | --- |
| 1<sup>st</sup> repeat | $k$ (kJ/(mol*day)) | $-0.08 \pm 0.03$ | $-0.09 \pm 0.04$ | $-0.09 \pm 0.02$ | – |
| | $\Delta G_0$ (kJ/mol) | $-28.4 \pm 0.75$ | $-28.9 \pm 1.9$ | $-28.6 \pm 0.5$ | – |
| 2<sup>nd</sup> repeat | $k$ (kJ/(mol*day)) | $-0.03 \pm 0.03$ | $-0.03 \pm 0.02$ | $-0.10 \pm 0.02$ | $-0.08 \pm 0.03$ |
| | $\Delta G_0$ (kJ/mol) | $-32.9 \pm 1.0$ | $-38.9 \pm 1.1$ | $-30.5 \pm 0.6$ | $-26.0 \pm 1.5$ |

**Table 2:** Experimental conditions used for fibril preparation in this study. The buffer composition and naming conventions were adapted _from_ [33,61–63].

| Condition | Buffer composition |
| --- | --- |
| FM | 50 mM Tris-HCl, 150 mM KCl, 1 mM EDTA, 0.05% NaN<sub>3</sub>, pH 7.5 |
| Ri | 5 mM Tris-HCl, 1 mM EDTA, 0.05% NaN<sub>3</sub>, pH 7.5 |
| F65 | 20 mM MES, 150 mM NaCl, 1 mM EDTA, 0.05% NaN<sub>3</sub>, pH 6.5 |
| F91 | 20 mM KPO<sub>4</sub>, 1 mM EDTA, 0.05% NaN<sub>3</sub>, pH 9.1 |

**Figure 1.**
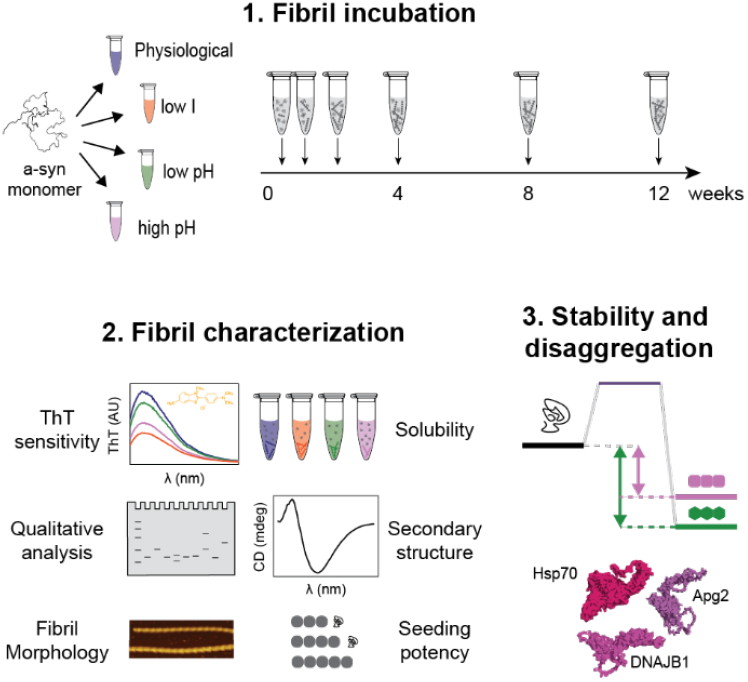
Overview of the experimental design. αSyn monomer was brought to four different sets of solution conditions – physiological (Fm, blue), low salt (Ri, orange), mildly acidic (‘low’) pH (F65, green), and mildly basic (‘high’) pH (F91, violet). Sample aliquots (2 × 0.5 mL, 100 μM for each timepoint and condition) were incubated at 37 °C and 600 rpm for 3, 6, 14, 28, 56, and 84 days. All samples were analyzed simultaneously by the indicated assays described in detail in the materials and methods section. The model structures of Hsp70 (P11142), DNAJB1 (P25685), and Apg2 (P34932) were taken from AlphaFold Protein Structure Database.^[67,68]^

### Time-dependent changes of αSyn polymorphism

First, we removed the supernatant from all samples by centrifugation and quantified the residual monomer concentration by UV-absorbance (Figure 2). Fm and F65 aggregated rapidly within two weeks of incubation whereas aggregation of Ri and F91 took approximately four weeks to reach a plateau. The monomer concentration converged to approximately 20 μM for all conditions except F91, for which it was significantly higher (60 μM). To rule out the possibility of a pseudo-equilibrium state caused by higher-order assembly of fibrils (observed in most of the tubes), we sonicated aliquots from the samples after 84 days of incubation to break down the fibril clumps and generate more seeding-competent fibril ends, and then extended the incubation period by five days. The monomer concentration in the supernatants of the centrifuged samples after this period did not change significantly in any condition (cross marks in Figure 2). The soluble species were not significantly degraded according to the SDS-PAGE analysis, and subsequent MS analysis revealed no substantial chemical modifications (Supplementary Figure 1, Supplementary Figure 2).

**Figure 2.**
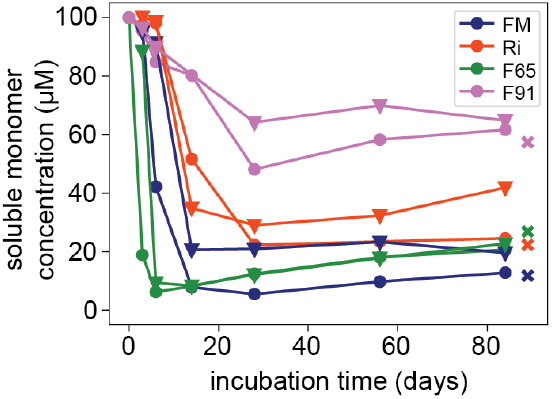
Residual monomer analysis. The individual points correspond to the monomer concentration in the supernatant after centrifugation at 16,000 x g at 25 °C for 90 min, determined by UV absorbance. The monomer concentration from 84-day samples which were sonicated and further incubated for five more days are indicated by the cross symbols. Round markers are data from the first repeat and triangles represent the second repeat (frozen and thawed).

After removing the soluble fractions, we analyzed the morphology, secondary structure content, and amyloid dye sensitivity of the fibrils using AFM, CD spectroscopy, and ThT assays, respectively. The CD spectra of sonicated fibrils showed a clear transition from a disordered monomer state to β-sheet rich species under all conditions (Supplementary Figure 3a). To assess the global variance across different time points and conditions, we performed a principal component analysis (PCA) on the CD spectra of all samples (Supplementary Figure 3b). The plot of the first two principal components, which together explain more than 95% of the variance, shows that the spectra of Ri and F91 form two distinct clusters, while those of Fm and F65 group together, suggesting structural similarity between the latter two (Supplementary Figure 3b). Interestingly, slight changes were observed in the CD spectra of samples from the second repeat, which had been frozen and thawed prior to measurement. The effect of freeze-thawing on αSyn fibril structure and morphology has been previously observed and described.^[69,70]^ We further examined the time-evolution of CD spectra for each fibril condition using singular value decomposition (SVD). The variance could be primarily attributed to the changes in the second spectral component with a maximum at 207 nm and a minimum at 220 nm, characteristic for a mixture of β-sheets and disordered regions (e.g., turns, random coils) (Supplementary Figure 3c).^[71]^ The component changed monotonically with time for F91, and to a lesser degree for Ri. In contrast, biphasic and triphasic behaviours were observed for the Fm and F65 samples, respectively, suggesting more pronounced structural rearrangements during early stages of their maturation (3 to 28 days, Supplementary Figure 3c). This observation is further supported by the sensitivity of the Fm samples to ThT, which shifted from high (days 3 and 6) to low (day 14 and older, Supplementary Figure 4a) and their seeding efficiency (Supplementary Figure 4b, Supplementary Figure 5). F65 samples displayed an erratic and nonlinear dependence of ThT fluorescence on fibril concentration, unlike F91 and Ri, which exhibited a linear relationship that remained consistent during their aging (Supplementary Figure 4a). Altogether we conclude, that the Fm and F65 appear structurally similar and undergo more significant changes during their aging compared to the Ri and F91.

Notable changes in fibril morphology during maturation were also observed in Fm and F65 samples through AFM analysis (Figure 3). In the Fm condition, early samples (3 days) contained thin fibrils (3-5 nm in height) with no detectable twist, which bundled into thicker filaments (Figure 3a). By day 6, the samples were predominantly composed of twisted fibrils with a pitch of around 200 nm and a height of 8 nm. A smaller population of shorter twisted fibrils (with a 150 nm pitch) was also present, and their numbers increased over the following week (up to 14 days). These shorter-twisted fibrils persisted after 84 days of incubation, albeit with a reduced height of around 5 nm, suggesting either tighter packing or truncation. However, SDS-PAGE analysis did not show any signs of the latter (Supplementary Figure 1 a, b). In the low pH (F65) condition, a well-defined population of 7-8 nm thick fibrils with a 150 nm pitch formed within 3 days and gradually matured into thicker fibrils (8-9 nm) with a slightly longer twist (about 170 nm, Figure 3b). Polymorphism within a given condition was most apparent between days 3 and 14, where minor populations of fibrils with distinct (or lack of) twists were observed. The polymorphic landscape of αSyn fibrils appeared less rugged for the Ri and F91 conditions (Supplementary Figure 6). In the low-salt samples (Ri), long, untwisted filaments, 6-7 nm thick, were observed after 14 and 84 days of incubation (Supplementary Figure 6 b). Similarly, fibrils formed in the high pH condition (F91) lacked any detectable twist (Supplementary Figure 6 a). These were initially thinner (around 4 nm) compared to those in Ri but gradually matured into 5-6 nm thick fibrils. In addition, the F91 samples contained a notable population of spherical aggregates of varying sizes, which may represent short fibril fragments or small amorphous aggregates.

**Figure 3.**
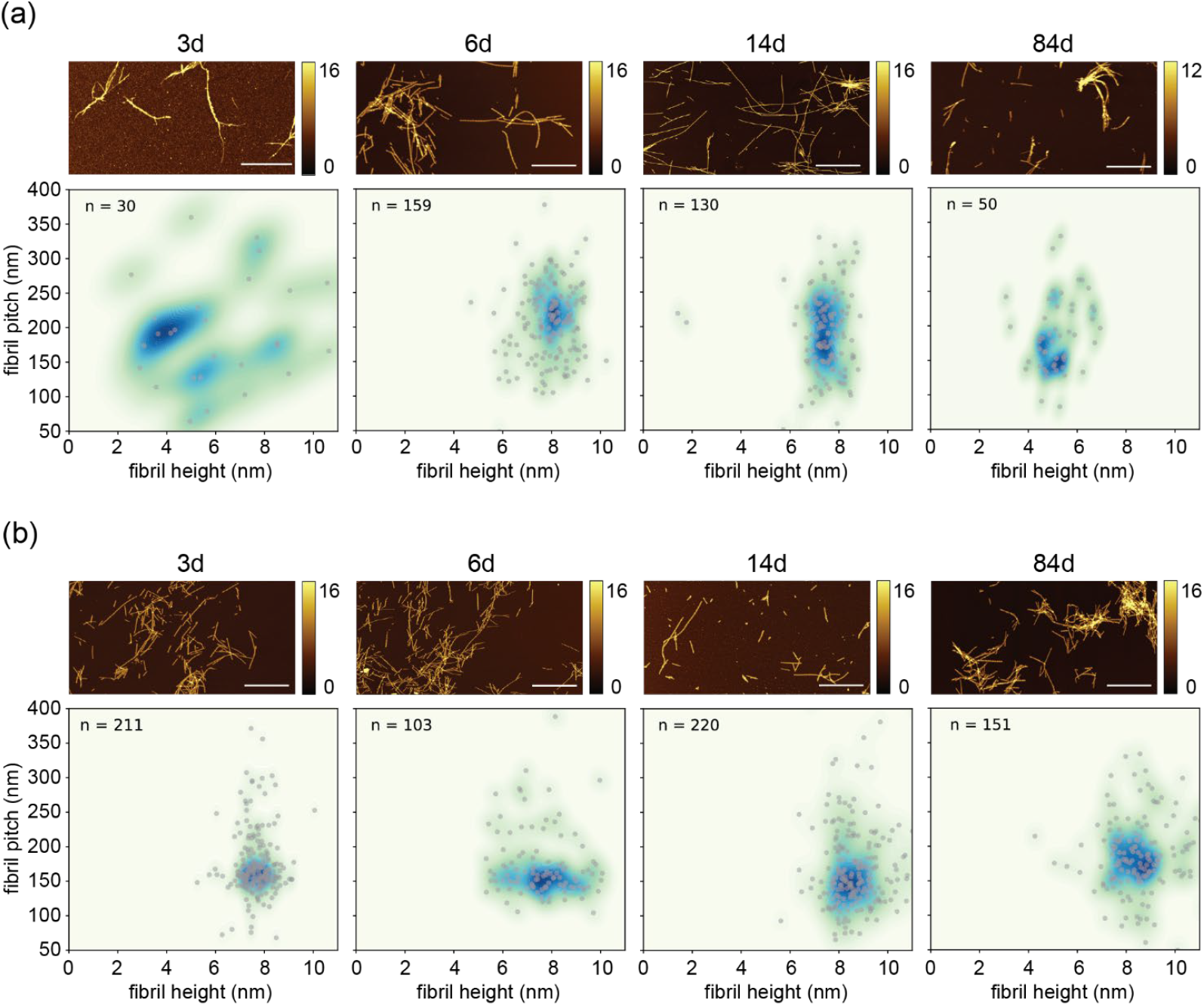
Time-dependent morphological changes of (a) Fm and (b) F65 fibrils monitored by AFM. Fibrils from four selected time points (3, 6, 14, and 84 days) were spotted on freshly cleaved mica and imaged using AFM tapping mode in air. At least two 20 μm × 20 μm independent spots were imaged for each sample with 10 nm/pixel resolution. Manually extracted fibril profiles were analyzed in terms of their pitch length (i.e., full 360° twist) and height using an automated python script (grey circles).^[43,72]^ Kernel density function (density gradient from light green to blue) was applied to each graph for better visualization of the fibril populations. The majority of Ri and F91 fibrils do not have twists and results of their AFM analysis are shown in Supplementary Figure 6. Scale bars represent 2 μm.

Based on our analyses and in accordance with previous reports, we conclude that fibrils formed under the four selected conditions exhibit distinct morphological characteristics. Moreover, we observe changes of polymorph composition during fibril maturation as described for αSyn,^[34,39]^ and other pathological amyloids such as Tau^[40]^ or IAPP.^[41]^

### Thermodynamic stability is an important determinant of fibril disaggregation by chaperones

Next, we used urea depolymerization to assess the thermodynamic stability of αSyn fibrils.^[43]^ The depolymerization was performed in the buffer used for chaperone disaggregation (50 mM HEPES, pH 7.5, 50 mM KCl, 5 mM MgCl2, 2 mM DTT) to allow its direct comparison to thermodynamic stability. The fibrils were centrifuged, the supernatant removed, and the fibril pellet resuspended and sonicated to achieve consistent concentration and homogeneity for all samples. The fraction of soluble monomer at increasing concentrations of urea was determined using flow-induced dispersion analysis (FIDA) as previously described^[43]^ and fitted using the isodesmic depolymerization model^[73]^. However, the stability (ΔG) parameter obtained by conventional fitting is correlated with the sensitivity towards denaturant (i.e., the m-value) which complicates the analysis of its time dependence ΔG(t). To overcome this and to better capture the relationship between the two fitting parameters, we employed Hamiltonian Monte Carlo (HMC) sampling of solutions and Bayesian analysis (Supplementary Figure 7, Supplementary Figure 8, Supplementary Figure 9, Supplementary Figure 10).^[74]^ First, we used HMC sampling of solutions to evaluate ΔG in an unbiased manner (Supplementary Figure 7a, Supplementary Figure 8a, Supplementary Figure 9a, Supplementary Figure 10a). We clearly observed an increase in stability of all four fibril types although still strongly correlated with the m-values (as expected). Fitting with a prior on the ΔG based on the solubility in the original conditions (Figure 2) failed to recapitulate the data, presumably because the stabilities of the different fibrils in the chaperone disaggregation buffer is different from their stabilities in the respective original buffers. Instead, we carried out the analysis with a moderate prior on the m-value (mean = 3.5 kJ mol^-1^ M^-1^, σ = 2, Supplementary Figure 7b, Supplementary Figure 8b, Supplementary Figure 9b, Supplementary Figure 10b)). The mean value was chosen based on the estimated changes of solvent accessible area between monomers in the amyloid and free state assuming it relates to the urea m-value analogously to what has been observed for unfolding of globular proteins (see Materials and Methods section for details of the calculation). The Bayesian analysis enabled more confident estimation of individual ΔG values whilst preserving the overall qualitative behaviour of ΔG(t) (Figure 4a,b, Supplementary Figure 7b, Supplementary Figure 8b, Supplementary Figure 9b, Supplementary Figure 10b)). The increase in stability was most prominent for F65 and F91, followed by Fm and Ri (Figure 4c). Interestingly, the ΔG of the Ri and Fm samples from the second repeat (grey triangles in Figure 4c) differed from the first repeat, suggesting structural or morphological alterations induced by the freezing and/or thawing as observed in the CD analysis (Supplementary Figure 3a).

**Figure 4.**
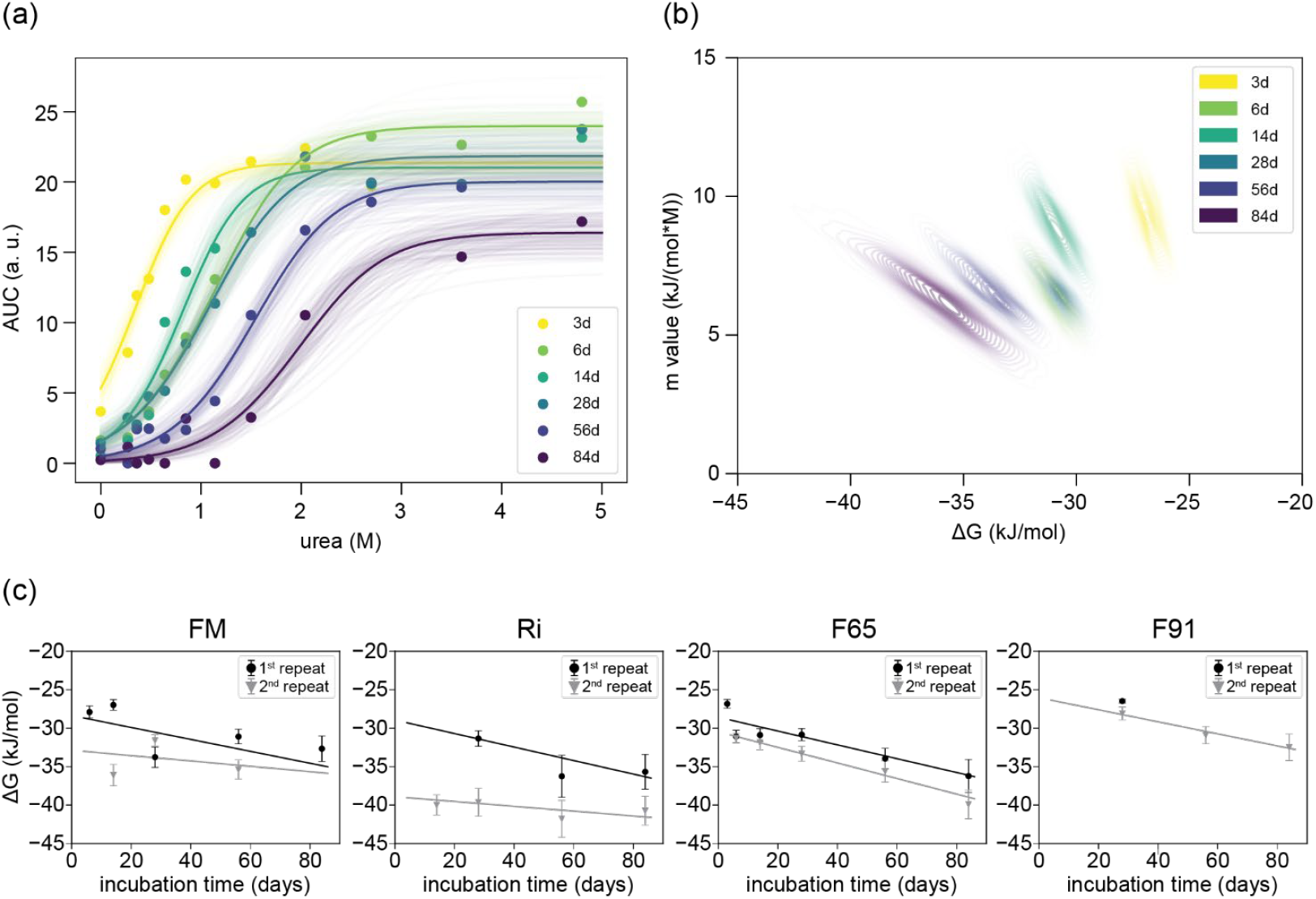
Thermodynamic stability of fibrils formed in conditions Fm, Ri, F65 and F91 and measured in 50 mM HEPES, pH 7.5, 50 mM KCl, 5 mM MgCl2, 2mM DTT. (a) Fitting of the isodesmic model to depolymerization curves of amyloid fibrils formed in condition F65 at different time points using Bayesian analysis with a prior of 3.5 and a standard deviation of 2 for the m-value. Opaque lines represent 100 random samples from a Hamiltonian Monte Carlo (HMC) analysis with 2000 samples in total. The area under the curve (AUC) corresponds to the monomer concentration which was measured by FIDA. (b) Probability distributions of ΔG and m-values obtained from the fitting in (a) highlight the correlation between these two fit parameters. (c) Time evolution of fibril thermodynamic stability ΔG(t) obtained by the urea depolymerization experiments. Depolymerization curves were fitted to the isodesmic model using Bayesian analysis as demonstrated in (a,b). Error bars indicate the uncertainty derived from a Markov Chain Monte Carlo (HMC) analysis (mean and standard deviation of 2000 samples). The global fits of the data to a linear model describing their maturation over time with rate k and the intercept at t=0 (ΔG0) is depicted by the black and grey lines for first and second repeat, respectively. Parameter values (ΔG and m-values) can be found in Supplementary Table 1, and Supplementary Table 2.

To further minimize errors from fitting individual depolymerization curves and compare differences in the fibril maturation rates, we used Bayesian analysis and globally fitted the datasets from each condition using an empirical linear model of aging according to equation 1.

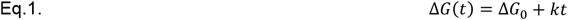

Here, ΔG_0_ is the hypothetical fibril stability at t = 0, and *k* is the rate constant of stabilization. The linear functions of ΔG(t) (Figure 4c, black and grey lines) agreed well with the values derived from individual fitting for all but one dataset (second repeat of Fm). The rates were significant (mean > 2SD) in all cases except the second repeats of Fm and Ri which we attribute to the alternations in fibrils structure induced by the freeze-thawing (Supplementary Figure 7, Supplementary Figure 8). The rates were comparable among the different fibril types and almost identical for F65 and F91. The ΔG0 values of Fm, F65 and Ri (Table 1) correspond to equilibrium monomer concentrations of ca 10 μM, whereas that of the F91 (2^nd^ repeat) ca 28 μM, reflecting the difference in solubilities measured in these conditions (Figure 2). Overall, our analysis revealed that Ri fibrils are the most stable, followed by Fm and F65, with F91 being the least stable.

Finally, we subjected the fibrils formed in the four different conditions to the chaperone system consisting of HSP70, DNAJB1 and Apg2 in the same buffer system (50 mM HEPES, pH 7.5, 50 mM KCl, 5 mM MgCl2, 2mM ATP, 2mM DTT), and used ThT to monitor their disaggregation (Figure 5).^[51,55]^ The fibrils were centrifuged, the supernatant removed, and the fibril pellet resuspended and sonicated to achieve consistent concentration and homogeneity across all conditions and time points. All fibril types were disaggregated with varying efficiencies except for Ri, as observed before (Figure 5, Supplementary Table 4).^[60]^ Interestingly, the ThT signal of Ri in the disaggregation assay increased in the presence of ATP (Supplementary Figure 12). This increase cannot be due to an increase in fibril concentration, as the remaining monomer was removed before the assay, as described above. We hypothesize that the observed increase in fluorescence may result from changes in ThT sensitivity, driven by the reduced lateral stacking of fibrils when chaperones bind in the presence of ATP. The disaggregation efficiency decreased with fibril maturation time for the other three fibril types (Figure 5b), suggesting that the structural rearrangement and/or increasing fibril stability impairs the chaperone activity. We observed that the thermodynamic stability correlates well with the degree of disaggregation (i.e., normalized change of the ThT signal before and after the reaction) for F65 and F91 fibrils (R^2^ of 0.85 and 0.70, respectively), and weakly for Fm (R^2^ of 0.25) (Figure 6). The slopes of correlations between stability and degree of disaggregation (Figure 6) observed for Fm, F65 and F91 reflect the magnitudes of the structural and morphological changes during their maturation. These are relatively minor in F91 fibrils which are all depolymerized to a similar degree (35-50%) despite the 5 kJ/mol stability difference between the early and the late ones. In contrast, the susceptibility to disaggregation varies significantly between early and late polymorphs of Fm (0-80%) and F65 (20-95%), reflecting their structural and morphological heterogeneity. The results clearly demonstrate that thermodynamic stability plays an important role but is not the sole determinant for fibril disaggregation by the chaperone system investigated here. A similar conclusion has been made in a recent study where neither stability nor affinity could fully explain the fibrils’ susceptibility for disaggregation.^[60]^ Here, we provide a new quantitative link between fibril stability and chaperone-mediated disaggregation, further highlighting the role of fibril morphology in determining the efficacy of chaperone action.

**Figure 5.**
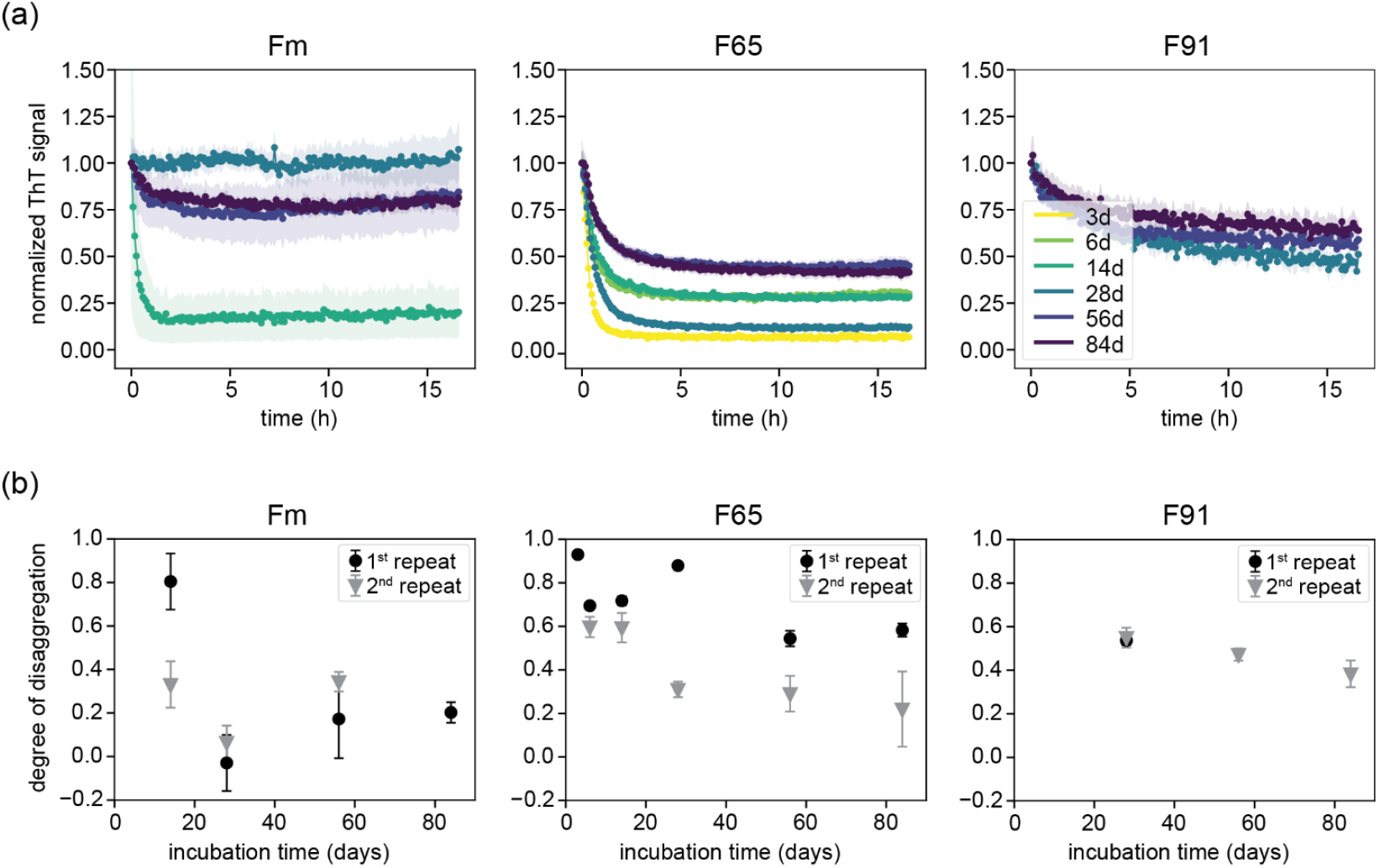
Kinetics of αSyn fibril disaggregation by the chaperone system HSP70, DNAJB1 and Apg2 in the presence of ATP. (a) Normalized ThT signal over time in the presence of chaperone system HSP70, DNAJB1 and Apg2, ATP and amyloid fibrils formed in either condition Fm, F65 or F91. Data corresponds to the first repeat in case of Fm and F65 and to the second repeat in case of F91. (b) Disaggregation efficiency of the chaperone system of amyloid fibrils formed in condition Fm, F65 or F91 at different time points of two repeats. A degree of disaggregation of 1 represents the complete loss of ThT signal. A degree of disaggregation of 0 represents no change of the ThT signal during the measurement. Error bars represent the propagated uncertainties based on the standard error of the mean from triplicate measurements. Fibrils formed in condition Ri were not disaggregated (Supplementary Figure 12).

**Figure 6.**
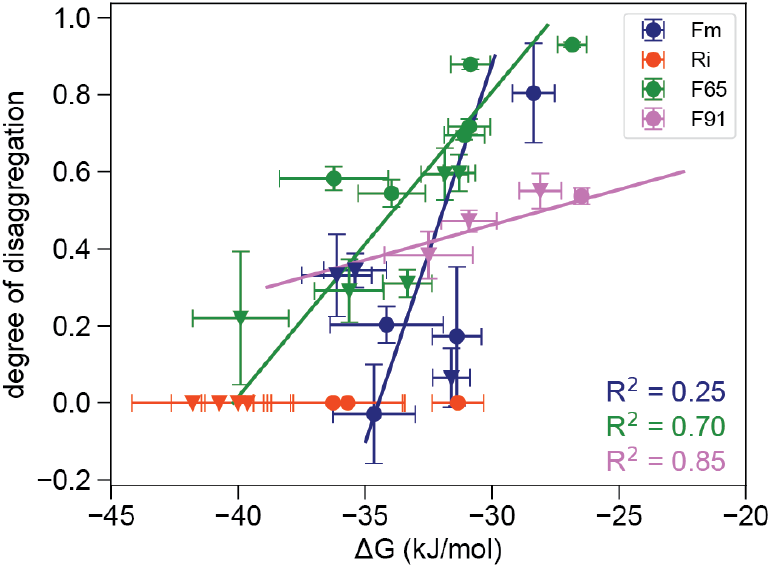
Correlation of the thermodynamic stability of αSyn fibrils and their disaggregation by the chaperone system HSP70, DNAJB1 and Apg2. Fibrils formed in condition Ri are not disaggregated. Their degree of disaggregation was set to 0 for comparison. The R^2^ value serves as a measure of the goodness of the linear fit to the data.

## Discussion and Conclusions

Here, we explored the relationship between amyloid fibril polymorphism, thermodynamic stability, and the susceptibility of αSyn fibrils to disaggregation by the HSP70/DNAJB1/Apg2 chaperone system. The structural features of fibrils change over time, and their thermodynamic stability increases. The changes and the increased stability may impact the ability of the chaperones to disaggregate fibrils, which could affect the onset of neurodegeneration. Therefore, this study was carried out to elucidate how the chaperone machinery reacts to time-induced changes in fibril morphology and thermodynamic stability. In line with earlier reports on αSyn and other amyloids, we observed significant shifts in the polymorphic distribution over extended timescales, well beyond the plateau phase of conventional ThT kinetic experiments.^[34]^ These shifts were evident through both bulk measurements such as altered ThT sensitivity, changes in secondary structure, and variations in seeding efficiency (Supplementary Figure 4, Supplementary Figure 5) as well as through single-particle analyses of fibril morphology by AFM, including height and pitch length (Figure 3, Supplementary Figure 6). These changes were most pronounced under conditions that favour rapid fibril formation, particularly in the presence of physiological salt concentrations (Fm) and low pH (F65). The coexistence and dynamic conversion of different fibril polymorphs in the Fm samples suggest that the αSyn amyloid landscape is the most rugged close to the neutral pH and physiological salt concentrations. At lower pH (F65), fibril formation is accelerated, favouring the emergence of kinetically accessible fibrils with similar morphologies and conformations, which continue to mature over longer timescales. In contrast, the energy barrier for fibril formation increases under low ionic strength conditions (Ri)^[33,43,75]^, arguably due to unfavourable electrostatic interactions. This limits the conformational space available for fibril formation and leads to highly stable fibrils.^[74]^ Similarly, at higher pH (F91), the fibril conformational space is reduced as the increased solubility of αSyn limits the number of energetically favourable fibrillar structures.

To our knowledge, this is the first quantitative description of the change in thermodynamic stability of αSyn fibrils formed under different conditions during maturation. The fibril stability ranged between ca -27 and -42 kJ/mol, consistent with previous studies.^[43,74,76]^ The Bayesian analysis of the chemical depolymerization allowed us to explore the relation between the ΔG and the m-values in detail. Interestingly, the average m-values obtained from fits of Ri and F91 depolymerization were close to the prior (ca 2 - 3.5 kJ mol^-1^ M^-1^, respectively) compared to the Fm and F65 which were significantly higher (ca 6 and 8 kJ mol^-1^ M^-1^, respectively). Notably, the m-values correlated weakly (R^2^ = 0.3) with AFM-derived fibril heights, suggesting that the m-values obtained from the isodesmic model may indeed reflect structural properties of the fibrils. The differences observed here may relate to distinct solvent exposure of the flexible N- and C-termini between the polymorphs, which were not accounted for when calculating the prior value.

Fibril maturation, represented by ΔG(t), was well described by an empirical linear model under all four conditions. The slow (∼months) fibril maturation occurs across all conditions without significant changes in the soluble monomer concentration after the first two to four weeks. Several potential mechanisms may be at play here: (i) slow structural rearrangements at the level of individual fibrils, (ii) Ostwald ripening, wherein less stable polymorphs gradually dissolve and “fuel” the growth of more stable ones or (iii) chemical modifications (oxidation, deamidation) of the protein, as well as degradation/truncation can influence the stability. Combination of high-resolution structural analysis and seeding experiments, supported by modelling of the reactions may help resolve these mechanisms. Furthermore, exploring the protein concentration dimension within the polymorphic landscape of αSyn fibrils in the future could elucidate the relative impact of nucleation, propagation and maturation of different polymorphs and their stability. Regarding point (iii) and based on our analysis of the protein by SDS-PAGE and mass spectrometry, chemical modification and truncation does occur to some extent but is not very pronounced and does not seem to affect a major fraction of the protein (Supplementary Figure 2). However, given that the majority of tyrosine residues is located in the C-terminus, even a modest degree of C-terminal truncation could lead to a misrepresentation of the soluble concentration at equilibrium, which does not decrease to values corresponding to our fitted ΔG values. However, our discovery of a clear correlation between fibril dissociation by a chemical denaturant and by chaperones is not affected by the exact manner in which the increase in apparent stability is generated.

Most importantly, our results corroborate previous observations regarding the relationship between fibril stability and disassembly of different αSyn polymorphs by the tri-chaperone system HSP70/DNAJB1/Apg2.^[60]^ We have found a clear correlation between ΔG and the degree of disaggregation for F65 and F91 and a weak correlation for Fm. This demonstrates that stability is a key factor in the disaggregation of amyloid fibrils by the tri-chaperone system. Fibrils formed in condition Ri are not disaggregated. Previous studies demonstrated that the lack of Ri fibril disaggregation cannot be explained by a weak chaperone binding and attributed this observation to their high stability inferred from cold denaturation.^[60]^ However, we see that the absolute stability of Ri is comparable to the three month sample of F65, which gets disaggregated. This clearly shows that fibril stability is indeed an important determinant of their disaggregation, but other factors must be at play. The difference in slopes of correlations in Figure 6 between individual fibril types are a strong indicator that polymorph-specific structural or morphological features are crucial for efficient chaperone action. These may include differences in the exposure of fibril core, packing, and/or dynamics of the flexible termini (i.e., fuzzy coat). The fuzzy coat is especially important as it contains chaperone binding sites ^[51,77,78]^. Different patterns of fibril lateral association or higher order assembly (e.g., fibril clumping) between the polymorphs might also affect the availability of the fibril surface or fibril ends.^[54]^ Further studies involving cryo-EM, hydrogen-deuterium exchange mass spectrometry coupled with crosslinking and other structural techniques are needed to resolve these factors in greater detail.

Importantly, our results demonstrate that the human tri-chaperone system can disaggregate fibrils as stable as -40 kJ/mol in certain cases. This value is significantly less compared to the free energy of ATP hydrolysis at the beginning of our depolymerization assays (ΔG = -68.5 kJ/mol, assuming 1% of initially hydrolyzed ATP, Figure 7).^[79]^ However, the value drops to -40.3 kJ/mol when 90 % of ATP is consumed and would recover only marginally (−46.3 kJ/mol) upon addition of fresh 2 mM ATP to the reaction. This suggests that the extent of ATP depletion, along with the accumulation of ADP and phosphate may set a stability threshold for fibrils that can still be disaggregated by the chaperones.

**Figure 7.**
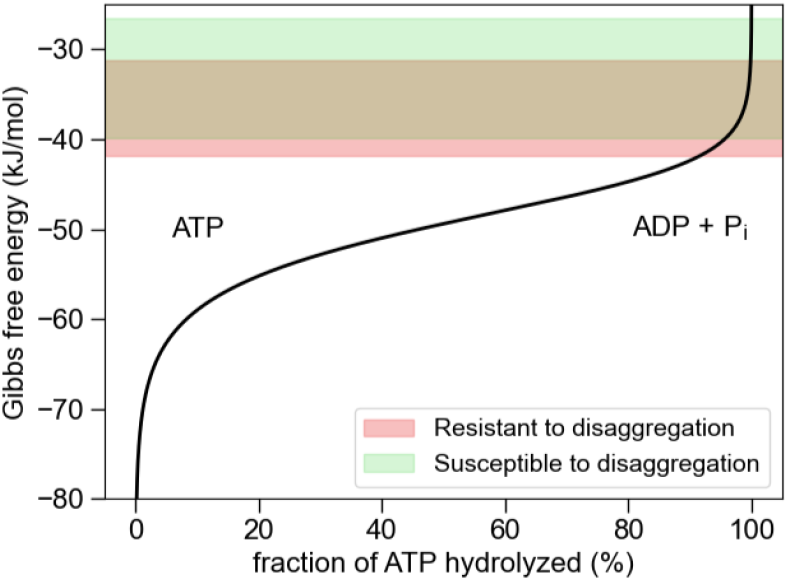
Change in the free energy of ATP as a function of its depletion. The Gibbs free energy of ATP hydrolysis to ADP and phosphate (P) (ΔG, black line) was calculated for conditions of the disaggregation assay (150 mM ionic strength, pH 7.5, 5 mM Mg^2+^) using the standard values from ^[79]^. The range of free energy corresponding to the fibril stabilities measured here are highlighted in red and green.

This observation holds significant implications for amyloid fibrils formed by proteins involved in other neurodegenerative diseases, such as Aβ, Tau, or IAPP. In the brain, amyloid fibrils persist over many years suggesting a high thermodynamic stability, which might lead to their resistance against disaggregation by the natural chaperone systems. Systematic studies of fibril stability and chaperone-mediated disaggregation, like the approach described here, will help to generalize our conclusions to amyloids formed from diverse protein sequences. Understanding the factors that determine the disaggregation propensity is a crucial step in understanding the molecular mechanisms in Parkinson’s and other neurodegenerative diseases. Fine tuning of the chaperone system (e.g., by optimization of the nucleotide recycling) or increasing the susceptibility of amyloid fibrils to the chaperone system might be a promising tool to reduce the amyloid fibril load in the brain.

## Experimental details

### Protein expression and purification

The chaperones DNAJB1 and Apg2 were expressed and purified as described before.^[51]^

For the expression of the human constitutive, cytosolic isoform of Hsp70, Hsc70 (HSPA8), chemo-competent *E. coli* BL21 (DE3) (New England Biolabs) cells were transformed by pCA528_SUMO_Hsc70 using heat shock. The expression of 6xHis-SUMO-Hsc70 was induced by IPTG (1 mM final concentration) and carried out at 20 °C for approximately 16 h. Cells were harvested by centrifugation (6000 x g, 4 °C, 15 min), resuspended in working buffer (WB, 50 mM HEPES, pH 7.4, 150 mM KCl, 5 mM MgCl2, 5% glycerol, 10 mM imidazole, 1 mM DTT) containing 1 mM PMSF and commercial DNase (Benzonase Nuclease, Sigma Aldrich), and lysed by sonication (Qsonica Q500, Newtown, CT, USA). The lysed solution was centrifuged at 17,000 rpm for 45 min at 4 °C. The supernatant was added to Ni-NTA beads (Thermo Scientific, Waltham, MS, USA) equilibrated in WB. The solution was incubated with the beads for one hour at 4 °C. Subsequently, beads were washed with WB containing 20 mM imidazole and bound proteins eluted by WB with 250 mM imidazole. Fractions containing 6xHis-SUMO-Hsp70 were cleaved by a SUMO protease (SAE0067, Merck KGaA, Darmstadt, Germany) during ca 16 h dialysis into 25 mM HEPES, pH 8.0, 150 mM KCl, 5 mM MgCl2, 5% glycerol, 2 mM DTT at 4 °C. On the next day, the solution was added to NiNTA beads and unbound protein including HSP70 was eluted by WB containing 20 mM imidazole. Fractions containing HSP70 were loaded onto a HiTrap Q HP, 5 mL column pre-equilibrated in 50 mM HEPES, pH 8.0, 10 mM KCl, 5 mM MgCl2, 1 mM DTT, Proteins were eluted by a 0-70% gradient of 1M KCl. Fractions containing HSP70 were pooled, concentrated and loaded onto a Superdex 200 10/300 increase GL (Cytiva) pre-equilibrated in 50 mM HEPES, pH 8.0, 150 mM KCl, 5 mM MgCl2, 1 mM DTT. Fractions containing highly pure HSP70 were flash frozen and stored at -80 °C.

αSyn expression and purification were carried out from pT7-7 αSyn WT plasmid (RRID:Addgene_36046) as described previously.^[43]^ In short, harvested cells were lysed using sonication (Qsonica Q500, Newtown, CT, USA, 10 s on time, 30 s off time, 12 rounds with 40% amplitude), and cell-free lysate further processed by boiling for 20 minutes. The resulting solution was precipitated by saturated (NH4)2SO4 (4 mL per 1 mL of supernatant, 15 min at 4°C) and centrifuged (20 000×g, 20 min at 4 °C). The resulting pellet was resuspended in 7 mL of 25 mM Tris–HCl pH 7.7 with 1 mM DTT and dialyzed twice into the same buffer. Monomeric αSyn was purified by anion exchange chromatography (AEC) (HiTrap Q Hp 5 ml, GE Healthcare) using linear gradient of NaCl, followed by size exclusion chromatography (SEC) (HiLoad 16/600 Superdex 200 pg. column, Cytiva). The monomeric fraction of αSyn eluted in 10 mM of NaP buffer pH 7.4 was collected, flash-frozen in liquid nitrogen and stored at -80°C.

### Fibril assembly

Fibrils were prepared in four different conditions provided in Table 1. Purified αSyn monomer was buffer exchanged into the desired assembly buffer using 3-kDa cut-off spin column filter (Amicon® Ultra Centrifugal Filter, Merck). The concentration was adjusted to 100 μM and the sample in each solution condition split into 12 aliquots of 0.5 mL which were flash-frozen in liquid nitrogen and stored at -80 °C. Two aliquots from each condition were thawed and placed into the incubator (Eppendorf ThermoMixer C, Germany) at 37 °C at 600 rpm at six different time points such that their incubation periods (3, 6, 14, 28, 54, and 84 days) ended simultaneously. After 84 days, one set of aliquots per timepoint per condition (i.e., n=24) was used directly in the experiments described further below whilst the other was flash frozen and stored at - 80 °C (i.e., 2nd repeat) and used as a repeat.

### Analysis of residual monomer concentration

A 50 μL aliquot was taken from each sample and centrifuged (16,000 x g, 90 min, 25°C). The resulting supernatant was carefully transferred to a clean tube and analyzed using SDS-PAGE, flow-induced dispersion analysis (FIDA), and UV-absorbance. The latter was carried out using a NanoDrop instrument (ThermoFisher, USA) and the extinction coefficient of αSyn (ε_280_ = 5,960 cm^-1^M^-1^) calculated from the sequence using Expasy webserver.

### SDS-PAGE

Samples were mixed with NuPAGE™ LDS Sample Buffer (4X) (Invitrogen, Waltham, MA, USA), loaded to a 4-20% pre-casted polyacrylamide gel (Biorad, USA), and analyzed under constant voltage of 200 V for 35 minutes. Proteins in the gel were stained by 1-hour incubation in InstantStain Coomassie Stain (INST-1L-181, Kem-En-Tec Nordic A/S, Denmark), followed by overnight destaining in dH_2_O. The concentration of αSyn was determined by densitometric analysis of the gels by ChemiDoc Go (BioRad, USA) using the Image Lab software (BioRad, USA) from the samples of known αSyn concentrations run in parallel as calibration.

### Intact mass spectroscopy of depolymerized fibrils

Fibrils were spun down as described earlier and the supernatant was removed. Solution of 8 M urea in 10 mM NaP buffer, pH 7.4 was added to depolymerize fibrils and samples were incubated for 3h at room temperature. Subsequently, the buffer was exchanged to 10 mM NaP buffer, pH 7.4 to remove urea from the sample. Samples were analyzed on an Ultraflex II MALDI-TOF/TOF Matrix-Assisted Laser Desorption Ionization (MALDI) mass spectrometer (Bruker). Samples for intact mass analysis were prepared for spotting by mixing 0.5 μL of sample with 0.5 μL of α-cyano-4-hydroxy-trans-cinnamic acid (saturated solution in 70% acetonitrile, 0.1% TFA). In addition to external calibration, α-Syn (14460 Da) sample was used as an internal standard to improve accuracy.

### Flow-induced dispersion analysis

The oligomeric state analysis of the supernatant and monomer quantification were further determined using the FIDA1 instrument (FidaBio, Denmark). Samples were analyzed using the following method:

1. Wash 1 (1M NaOH): 45 s, 3500 mbar.
2. Wash 2 (MQ water): 45 s, 3500 mbar.
3. Equilibration (Buffer): 30 s, 3500 mbar.
4. Sample application (Protein): 20 s, 75 mbar.
5. Measurement and detection (Buffer): 75 s, 1500 mbar.

All FIDA experiments were performed at 25 °C. The monomer concentration was quantified from the areas under the peak (obtained by fitting a Gaussian function to the Taylorgrams) using a calibration curve of known monomer concentrations.

### Thioflavin T sensitivity assay

The sensitivity of the individual samples to Thioflavin T (ThT) was measured in 384-well low volume non-binding assay plates (Corning 3544) using a VANTAstar plate reader (BMG, Germany). A 1:1 dilution series of four samples was prepared for each aliquot using the respective buffer and supplemented with 50 μM ThT from a concentrated stock solution (5.5 mM). Each Fluorescence spectrum was measured in the 465-600 nm wavelength range upon excitation at 440 nm in 1 nm increments. The maximal intensity was plotted against fibril concentration calculated from the residual monomer analysis. Each sample was measured in duplicates (2 × 15 μL).

### Atomic force microscopy

All samples were diluted by the appropriate buffer to 5 µM monomer equivalent concentration (calculated by subtracting the residual monomer concentration from the total concentration) and 25 µL of the solution was deposited onto freshly cleaved mica substrates. Following 2 min of incubation, the substrates were extensively cleaned by miliQ water and dried under nitrogen gas flow. All fibrils were imaged in tapping mode in air using a DriveAFM (Nanosurf, Liestal, Switzerland) using PPP-NCLAuD cantilevers (Nanosensors, Neuchatel, Switzerland). The AFM images were processed using the Gwyddion software. The pitch length and height analysis of fibrils was carried out from the manually extracted fibril profiles using an automated python script. ^[43,72]^

### Fibril sonication

A 200 μL aliquot was taken from each sample and centrifuged (16,000 x g, 90 min). Most (180 μL) of the supernatant was carefully removed and the pellets resuspended in their respective buffer to a monomer-equivalent concentration of 200 μM. Resuspended fibrils were sonicated using an ultrasonic probe (Hielscher UP200St, Germany) in 3s-pulses of 20% amplitude separated by 12 s pauses for 5 minutes (one minute total sonication time).

### Circular dichroism

Sonicated fibrils were diluted to 15 μM and their circular dichroism (CD) spectra were recorded using a J-1500 CD spectrometer (Jasco, Japan), The spectra were collected in the 190-250 nm wavelength range at 50 nm/min scan rate and 0.5 nm steps at 25 °C. The final reported spectra are an average from triplicate measurements. The CD spectra of each buffer and αSyn monomer were collected as a reference.

### Thioflavin T seeding assay

The seeding potency of each fibril sample was measured in 384-well low volume non-binding assay plates (Corning 3544) using a FLUOstar plate reader (BMG, Germany). The monomer was diluted into disaggregation assay buffer (50 mM HEPES, pH 7.5, 50 mM KCl, 5 mM MgCl_2_, 2mM DTT) to final concentrations of 5, 10, 20, and 40 μM and supplemented with ThT (50 μM final concentration). Fixed volumes of sonicated fibrils (2.5 μM concentration, or the highest possible for the samples with low fibril concentration) were added to each sample and ThT fluorescence upon excitation at 440 nm was monitored at 480 nm under quiescent conditions in 5 min intervals at 37 °C. The initial parts of the curves (the first 2.5 hours) were fitted to a linear function, and the resulting slopes plotted against the initial monomer concentrations. The slope of the resulting curve corresponds to the apparent elongation rate (*k*^app^) which is a product of the elongation rate constant (*k*^+^) and concentration of seeding-competent ends.

### Urea depolymerization

Sonicated fibrils were diluted into 1x assay buffer (50 mM HEPES, pH 7.5, 50 mM KCl, 5 mM MgCl2, 2mM DTT) containing 0 to 4.8 M urea (40 µM final concentration in monomer-equivalents). Samples were incubated at room temperature for 3 days to allow for equilibration at the respective urea concentrations. After the incubation, the samples were analyzed by FIDA using the method described above. The monomer concentration was extracted from the elution profiles after correction for the viscosity at different urea concentrations as previously described.^[43]^ The chemical depolymerization curves were analyzed using NumPyro^[80]^ to sample posterior distributions of the isodesmic model (Equation 2)^[74]^ parameters using the No U-Turn Sampler (NUTS).

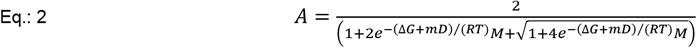

Where A is the area under the Gaussian peak from FIDA, M is the protein concentration in monomer equivalents, ΔG is the thermodynamic stability, m is the m-value, R the universal gas constant, and T is the temperature.

Data was analyzed using flat priors to visualize the maximum-likelihood estimation (MLE) or with a gaussian prior on the m-values (denaturant dependence of thermodynamic stability) for a Bayesian treatment. The prior was based on its relationship to changes in solvent accessible area (ΔSASA) between structured (i.e., fibrillar) and disordered state (i.e., monomeric), derived from analysis of globular protein unfolding.^[81]^ The SASA values for the αSyn monomers in the fibrillar states were calculated from 13 representative structures of αSyn fibrils (pdb IDs 2n0a, 6a6b, 6cu7, 6cu8, 6osm, 6sst, 6ssx, 7nck, 7yk2, 7ynm,8bqv, 8pk2, 9euu) using the FreeSASA python module.^[82]^ Only the SASA of a single monomer chain in the outermost layer of the fibrils was considered. The SASA of soluble, disordered state of αSyn monomers was calculated as a sum of per residue SASA (rSASA) contributions of residues buried in the corresponding fibril cores using average of different rSASA scales reviewed in ^[83,84]^.

### Chaperone disaggregation

α-Synuclein fibrils were disaggregated by the human chaperone system Hsp70+DNAJB1+Apg2.^[51]^ The fibrils were centrifuged at 16000 xg at 25 °C for 90 min in a tabletop centrifuge. The supernatant was discarded. The pellet was resuspended in the respective fibril buffer to a monomer-equivalent concentration of 200 µM. Fibrils were sonicated (1 min on, 1 min off, 1 min total sonication time, amplitude: 30%) using an ultrasonic probe (Hielscher UP200St, Teltow, Germany). Fibrils were disaggregated at 2 µM by 4 µM Hsp70, 2 µM DNAJB1 and 0.2 µM Apg2 in 50 mM HEPES, pH 7.5, 50 mM KCl, 5 mM MgCl_2_, 2 mM DTT and 2 mM ATP at 30 °C. The disaggregation was followed by 20 µM ThT with a VANTAstar plate reader (BMG LABTECH, Ortenberg, Germany) with a 440-10 nm excitation and 480-10 nm emission filter. Samples were measured every 5 min with orbital shaking for 3 s before each cycle in a 384-well plate (Corning 3544).

## Authors contributions

A.K.B. supervised the work. C.F., A.K., A.S.W., B.B. and A.K.B. conceptualized the work. C.F. and A.K. designed and carried out the experiments, analyzed the data, prepared the graphics, and wrote the manuscript. S.W. performed the mass spectrometry experiments. R.K.N. and C.F. wrote the python code for the analysis of urea depolymerization experiments. T.L.D. guided chaperone mediated disaggregation experiments. All authors contributed to the preparation of the manuscript and agree with its content.

## Acknowledgements

A.K.B thanks the Novo Nordisk Foundation for funding (NNF17SA0028392 and NNF21OC0065495). This research was co-funded by the European Union (ERC CoG 101088163 EMMA to A.K.B.), Lundbeck foundation (grant number R366-2021-169 STADIC to A.K.B.), and the Deutsche Forschungsgemeinschaft (DFG, German Research Foundation) – project number 504257241 to BB. A.K. would like to acknowledge support through a Horizon MSCA individual postdoctoral fellowship (Grant number 101106115) for funding.

