## Supporting information for "Thermodynamic Stability Modulates Chaperone-Mediated Disaggregation of α-Synuclein Fibrils"

### Supporting Figures

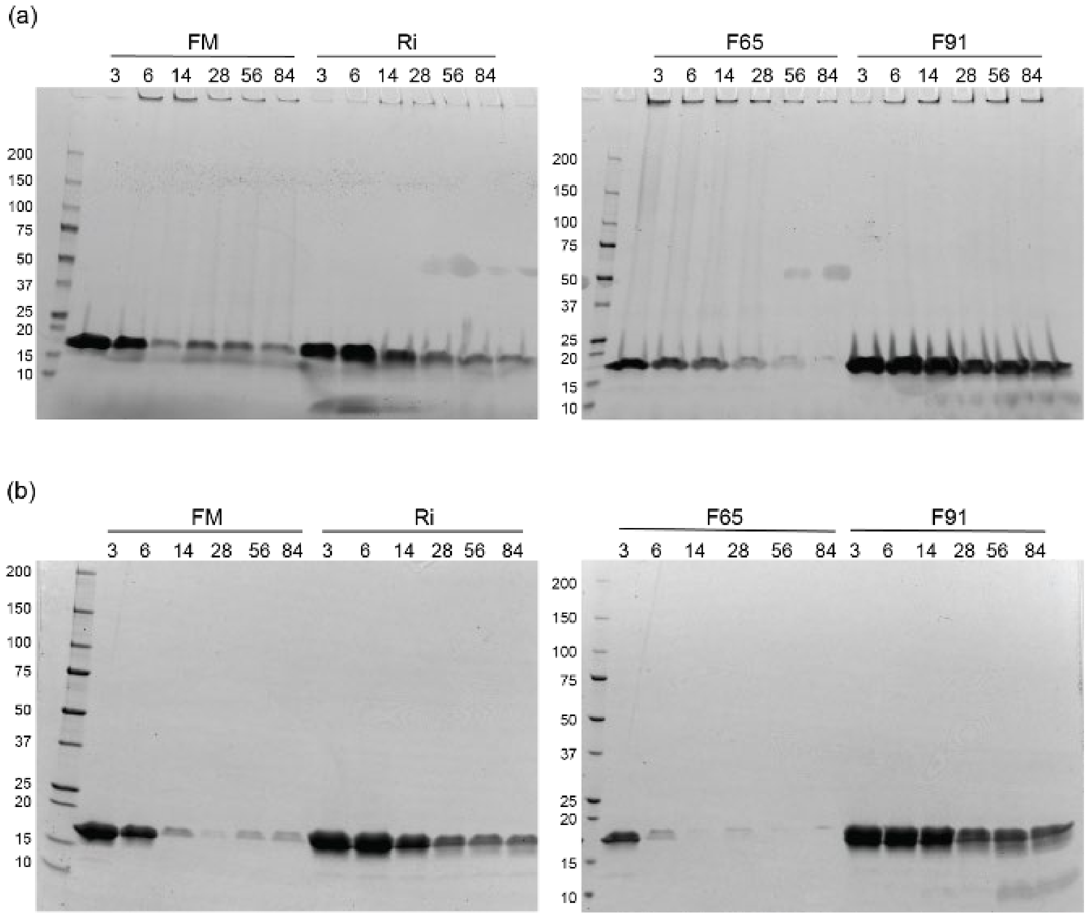


**Supplementary Figure 1.** SDS-PAGE of the whole sample (a) and the supernatant (b) of 100 µM αSyn incubating at 37 °C, 600 rpm for the indicated number of days. Shown samples are from the first repeat. The marker indicates the molecular weights of standard proteins in kDa. Samples for supernatant analysis were spun down at 16000 xg for 90 min. The supernatant was loaded into an SDS-PAGE gel.

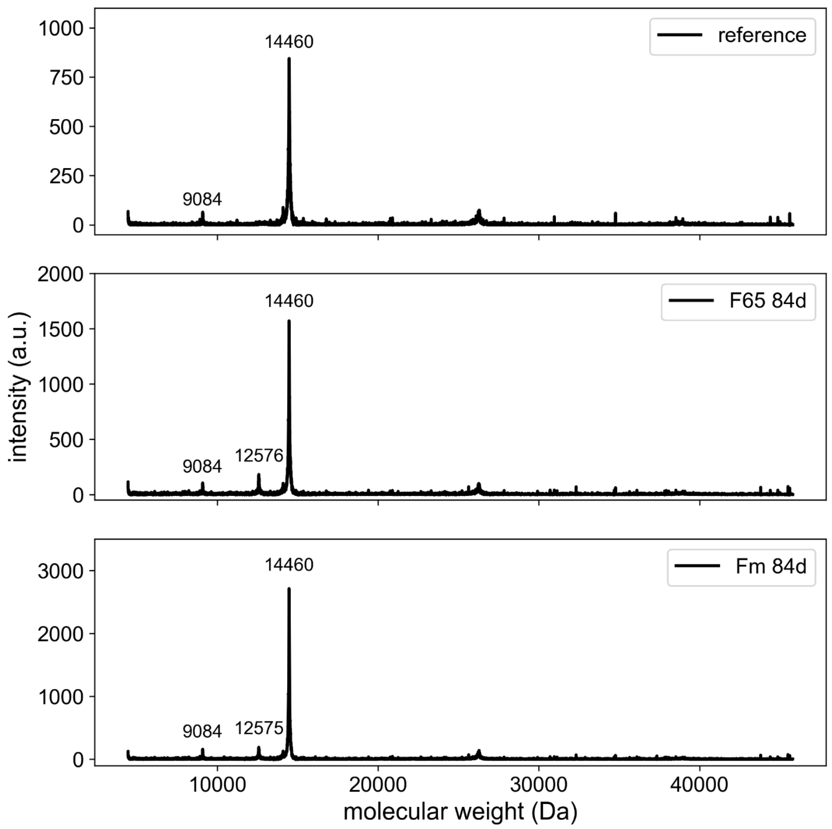


**Supplementary Figure 2**. Intact mass spectrometry of depolymerized fibrils of conditions F65 and Fm, with SEC purified, freshly thawed monomer for reference. The masses of the detected fragments are indicated for the corresponding peaks.

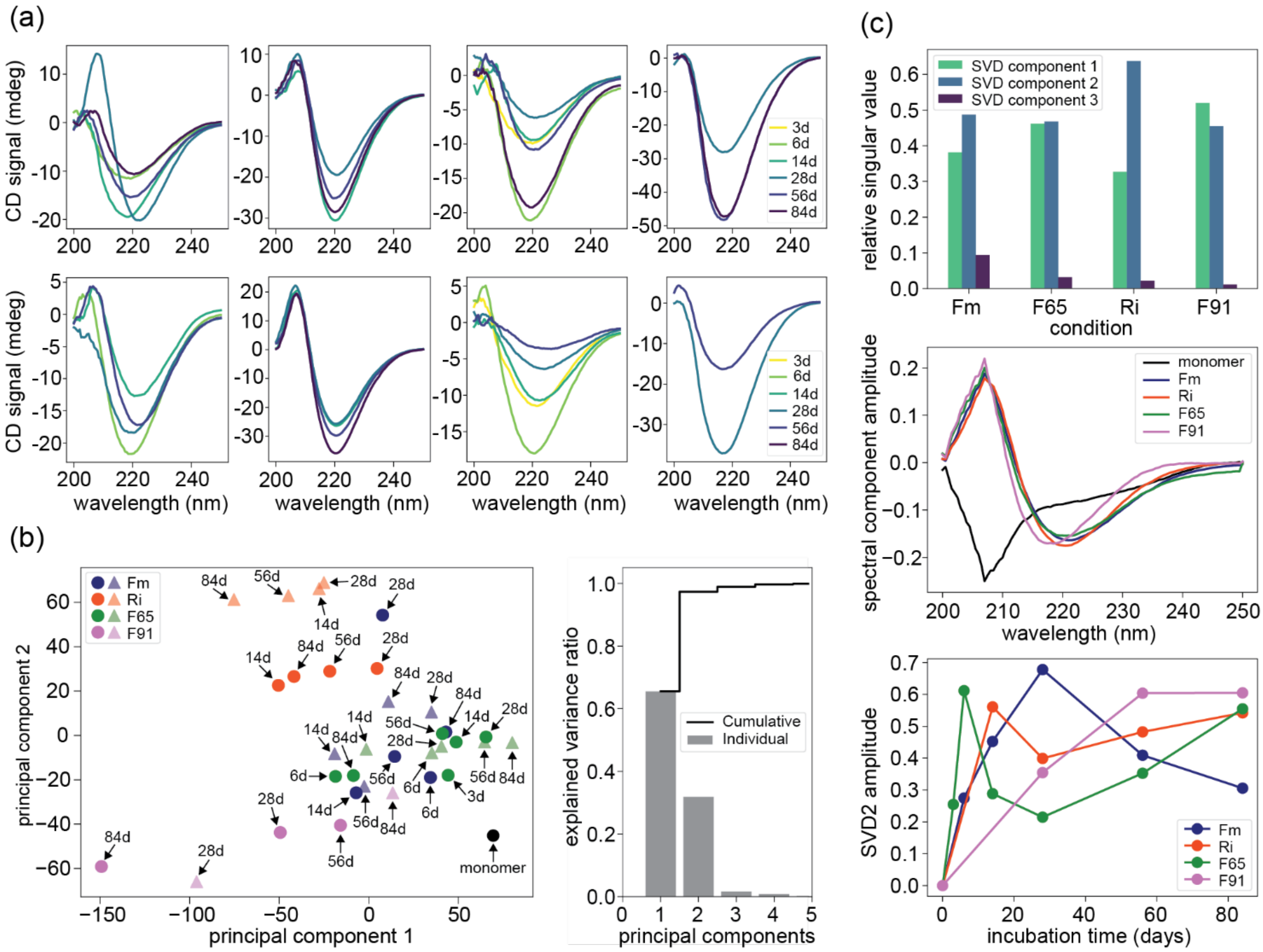


**Supplementary Figure 3: Circular dichroism analysis of the fibril maturation process.** (a) CD spectra of amyloid fibrils formed by αSyn in different conditions and at different time points. Upper panel: 1st repeat, lower panel: 2nd repeat. From left to right: Fm, Ri, F65, F91. Fibrils were centrifuged at 16,000 x g for 90 min. The supernatant was removed and the pellet containing fibrils was resuspended in the respective buffer condition. Samples were measured at 15 μM monomer equivalent concentration. (b) PCA analysis of all spectra shown in (a). left: principal component 1 vs 2. Dots represent the 1st repeat, triangles the 2nd repeat. Individual and cumulative variance explained by the components (right). (c) SVD analyses of the time-dependent changes in CD spectra in each condition shown in the upper panel of (a) (1st repeat). Top: Relative singular values of the first three SVD components. Middle: Spectral components obtained from the SVD analyses. The first component was fixed to the spectrum of the monomeric αSyn (black) for all conditions as a reference.[1] The second spectral component from each condition is color coded according to the legend. Bottom: Changes of second SVD component over time for αSyn fibrils formed in different conditions.

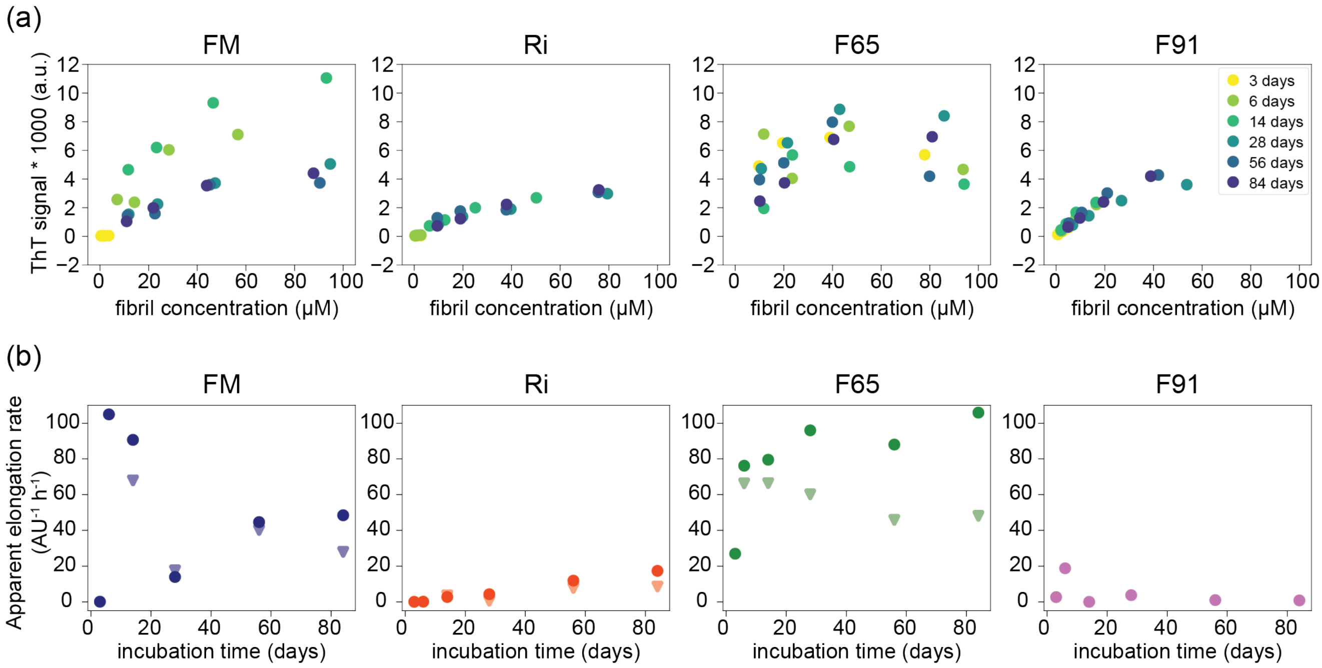


**Supplementary Figure 4:** ThT analysis and seeding efficiency of polymorphs of the 1st repeat at different time points. (a) ThT sensitivity of the fibrils. Concentration series of αSyn solutions at each time point were mixed with ThT and measured in plate reader. The concentration of fibrils corresponding to monomer equivalents were back-calculated from the soluble monomer concentration at each time point (figure 2). (b) Seeding efficiency of 2.5 μM sonicated fibrils (monomer equivalent), or the highest possible concentration for the samples with low fibril concentration, in 50 mM HEPES, pH 7.5, 50 mM KCl, 5 mM $MgCl_2$, 2mM DTT. Dots represent the 1st repeat, triangles the 2nd repeat.

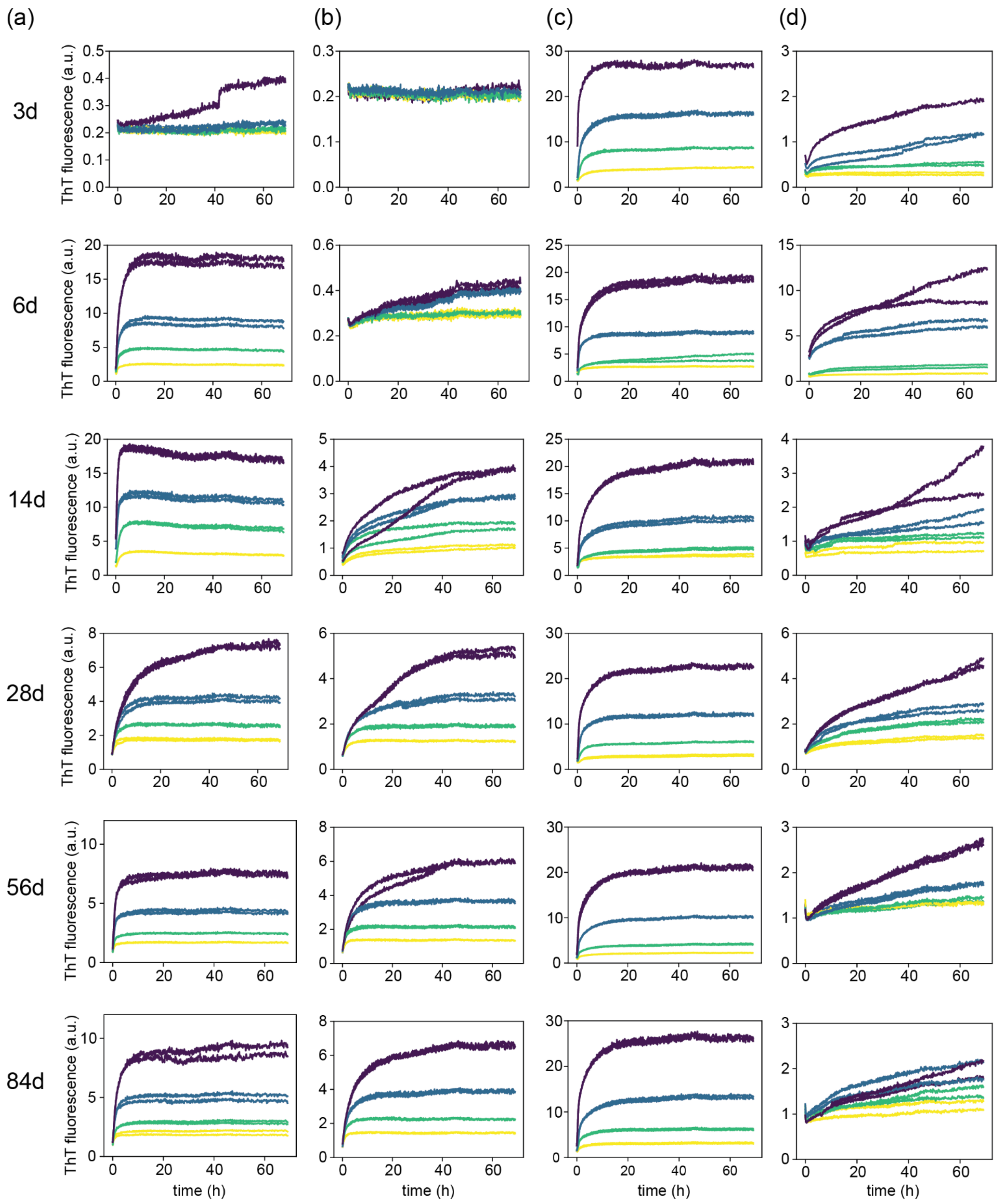


**Supplementary Figure 5.** Aggregation kinetics of αSyn monomer seeded by fibrils formed in formed in (a) Fm, (b) Ri, (c) F65, (d) F91 over time. The 2.5 μM seeds (i.e., sonicated fibrils) were added to 5 (yellow), 10 (green), 20 (blue), and 40 μM (purple) monomer solution in the disaggregation buffer (50 mM HEPES, pH 7.5 50 mM KCl, 5 mM $MgCl_2$, 2 mM DTT) supplemented with 50 μM ThT.

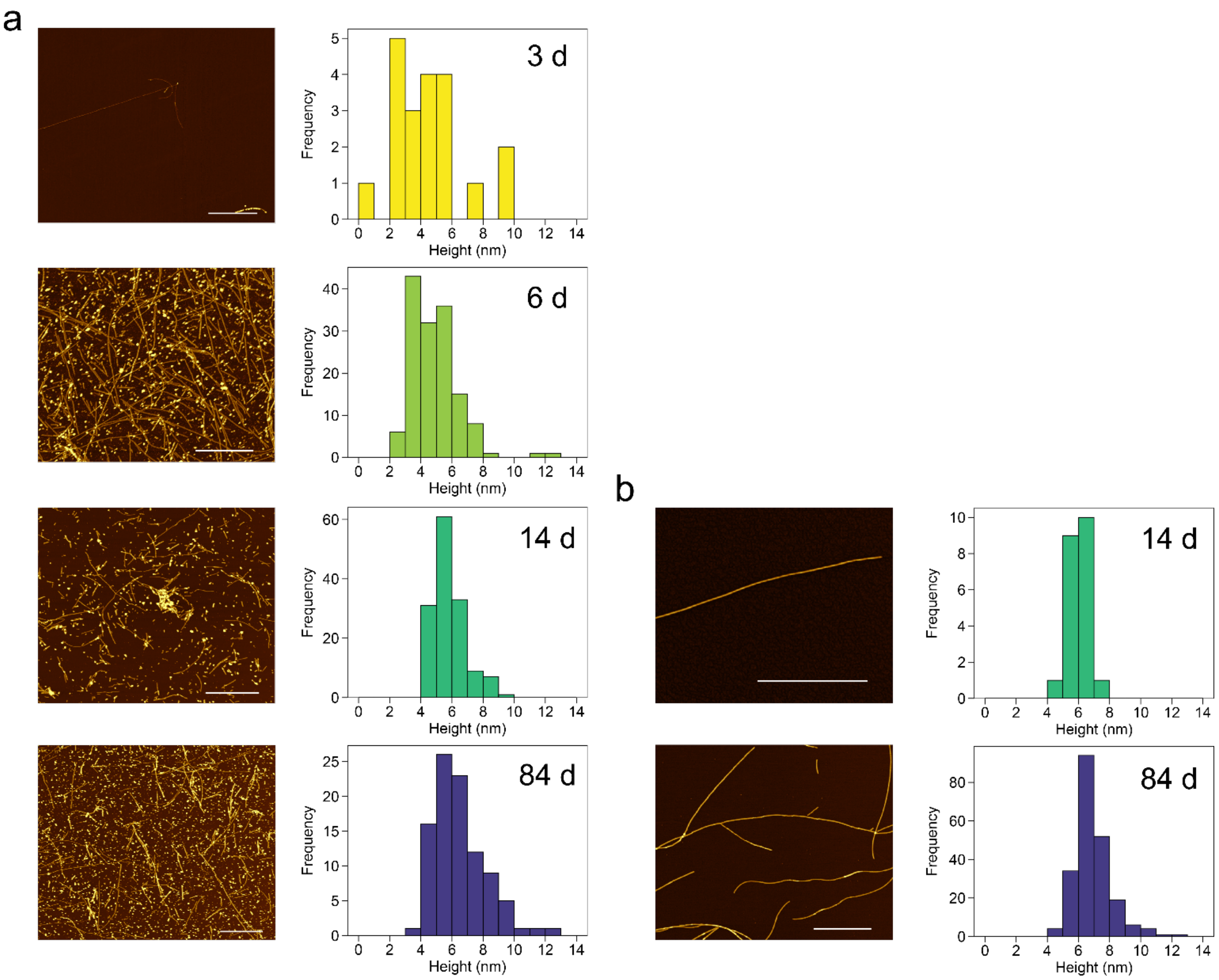


**Supplementary Figure 6.** Time-dependent morphological changes of (a) F91 and (b) FRi fibrils monitored by AFM. Fibrils from four selected time points (3, 6, 14, and 84 days) were spotted on freshly cleaved mica and imaged using AFM tapping mode *in air*. For Ri, no fibrils could be found after 3 and 6 days of incubation. At least two 20 µm x 20 µm independent areas were imaged for each sample with 10 nm/pixel resolution. Manually extracted fibril profiles were analyzed in terms of their height using an automated python script (right panels). Scale bars correspond to 2 µm.

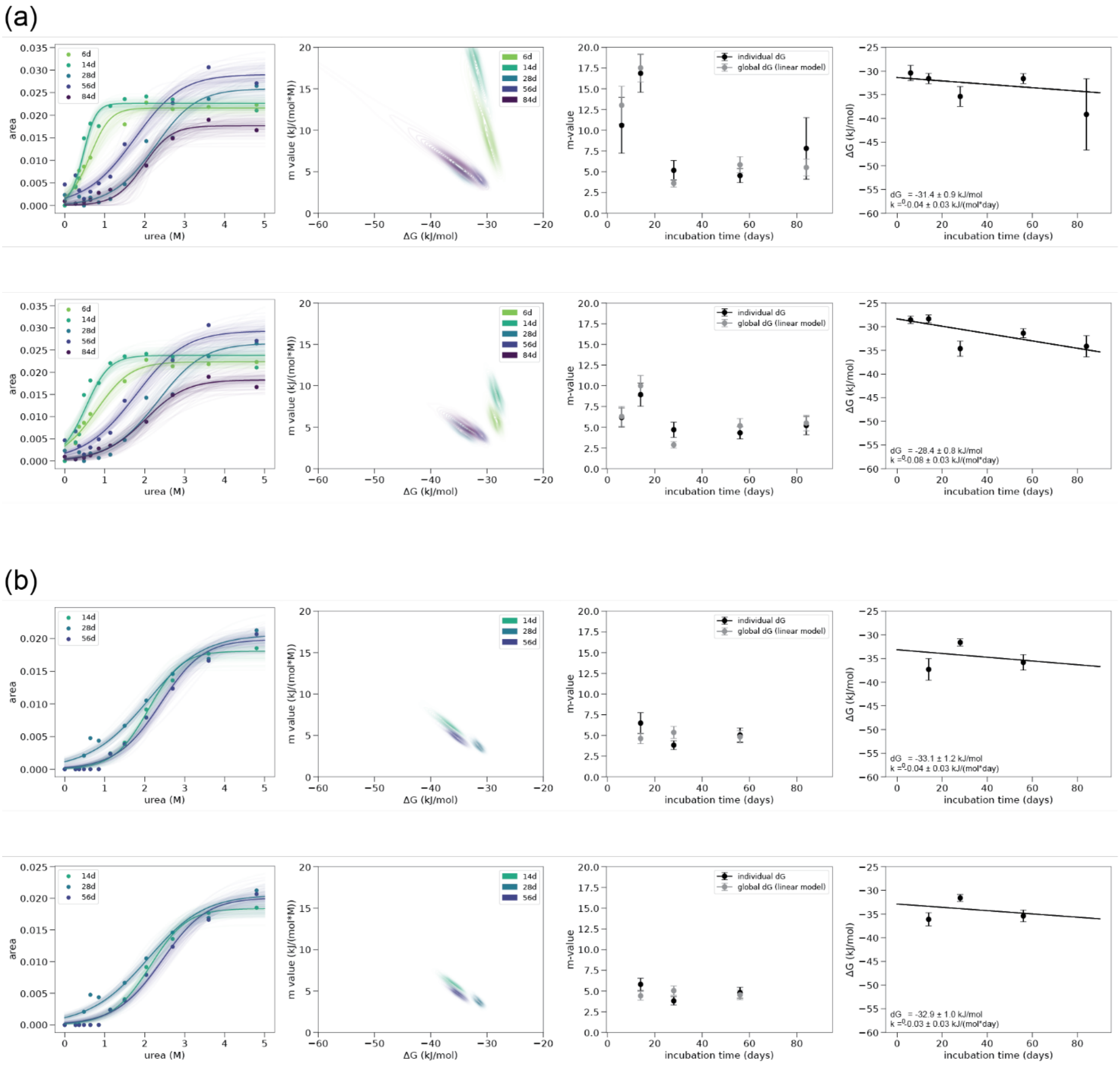


**Supplementary Figure 7.** Urea depolymerization curves of fibrils formed in condition Fm over time. (a) 1st repeat. (b) 2nd repeat. Curves were fitted using HMC sampling of solutions (n=2000) in the upper panel in (a,b) and using Bayesian analysis with a normally distributed prior on m of 3.5 with a standard deviation of 2 (lower panel in (a,b)). Column 1: Data points (area under the curve of FIDA measurements corresponding to monomeric αSyn) and their fits to the isodesmic model using HMC sampling of solutions or Bayesian analysis with HMC sampling of solutions. Column 2: Correlation of the m-value and ΔG. Column 3: Comparison of the m-value of the two different fitting approaches. Column 4: Fitted ΔG values over time (mean of 2000 solutions, error bars represent the standard deviation of the 2000 solutions) and fitting of the data to a linear model with decreasing ΔG over time: $\Delta G(t) = \Delta G_0 + k*t$. The urea depolymerization series was measured in 50 mM HEPES, pH 7.5 50 mM KCl, 5 mM $MgCl_2$, 2 mM DTT.

(a)

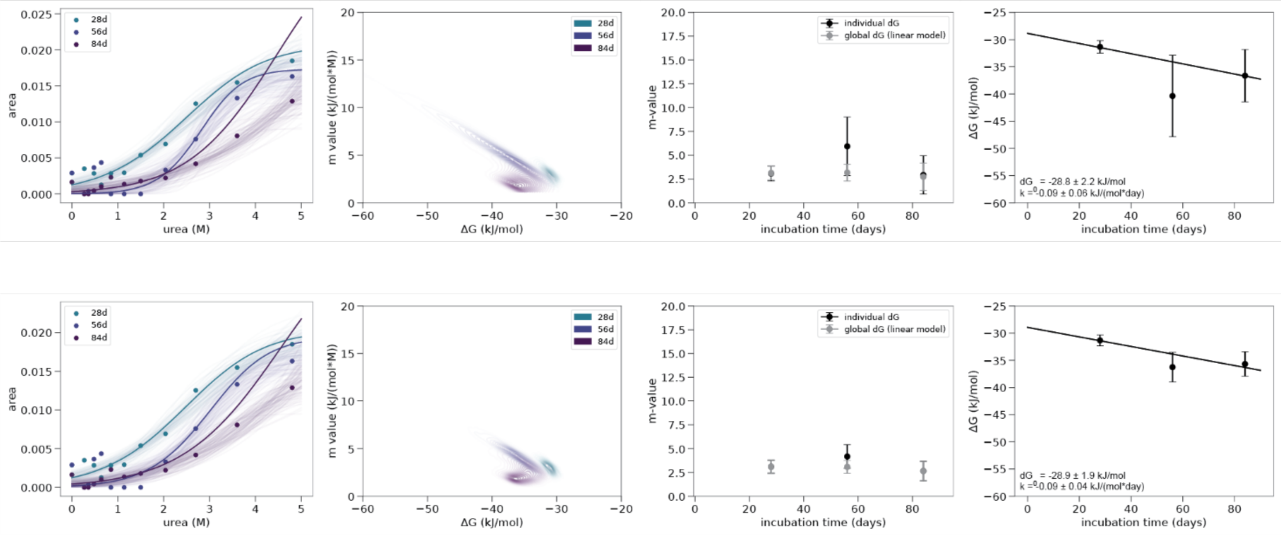


(b)

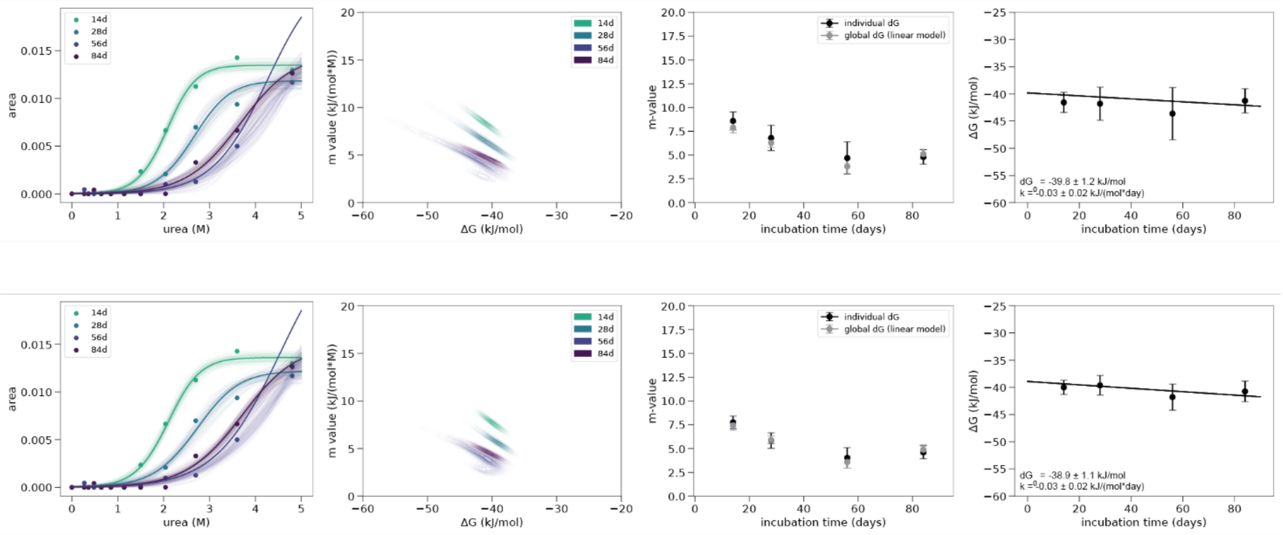


**Supplementary Figure 8.** Urea depolymerization curves of fibrils formed in condition Ri over time. (a) 1st repeat. (b) 2nd repeat. Curves were fitted using HMC sampling of solutions (n=2000) in the upper panel in (a,b) and using Bayesian analysis with a normally distributed prior on m of 3.5 with a standard deviation of 2 (lower panel in (a,b)). Column 1: Data points (area under the curve of FIDA measurements corresponding to monomeric αSyn) and their fits to the isodesmic model using HMC sampling of solutions or Bayesian analysis with HMC sampling of solutions. Column 2: Correlation of the m-value and ΔG. Column 3: Comparison of the m-value of the two different fitting approaches. Column 4: Fitted ΔG values over time (mean of 2000 solutions, error bars represent the standard deviation of the 2000 solutions) and fitting of the data to a linear model with decreasing ΔG over time: $\Delta G(t) = \Delta G_0 + k*t$. The urea depolymerization series was measured in 50 mM HEPES, pH 7.5 50 mM KCl, 5 mM $MgCl_2$, 2 mM DTT.

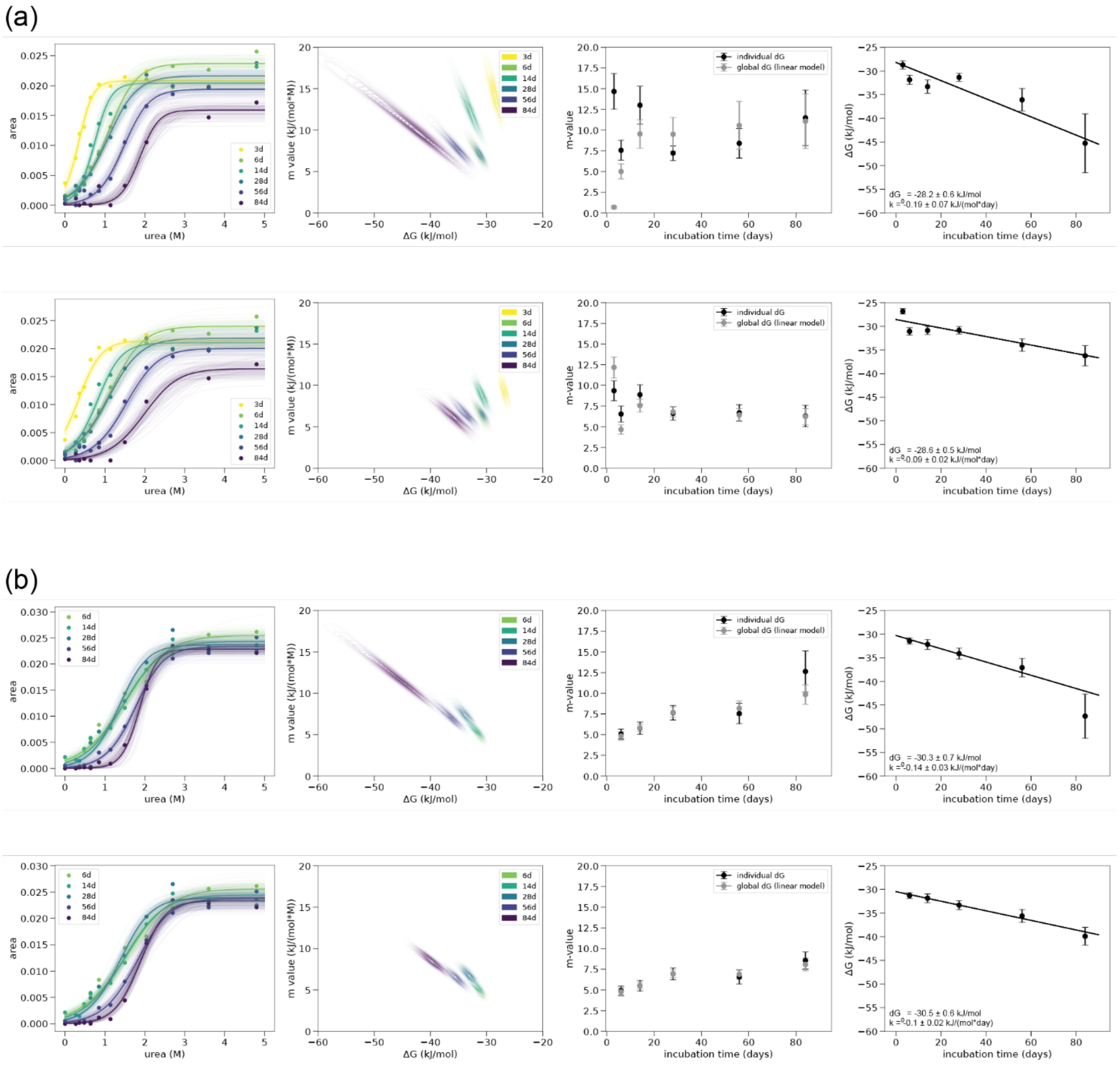


**Supplementary Figure 9.** Urea depolymerization curves of fibrils formed in condition F65 over time. (a) 1st repeat. (b) 2nd repeat. Curves were fitted using HMC sampling of solutions (n=2000) in the upper panel in (a,b) and using Bayesian analysis with a normally distributed prior on m of 3.5 with a standard deviation of 2 (lower panel in (a,b)). Column 1: Data points (area under the curve of FIDA measurements corresponding to monomeric αSyn) and their fits to the isodesmic model using HMC sampling of solutions or Bayesian analysis with HMC sampling of solutions. Column 2: Correlation of the m-value and ΔG. Column 3: Comparison of the m-value of the two different fitting approaches. Column 4: Fitted ΔG values over time (mean of 2000 solutions, error bars represent the standard deviation of the 2000 solutions) and fitting of the data to a linear model with decreasing ΔG over time: $\Delta G(t) = \Delta G_0 + k*t$. The urea depolymerization series was measured in 50 mM HEPES, pH 7.5 50 mM KCl, 5 mM $MgCl_2$, 2 mM DTT.

(a)

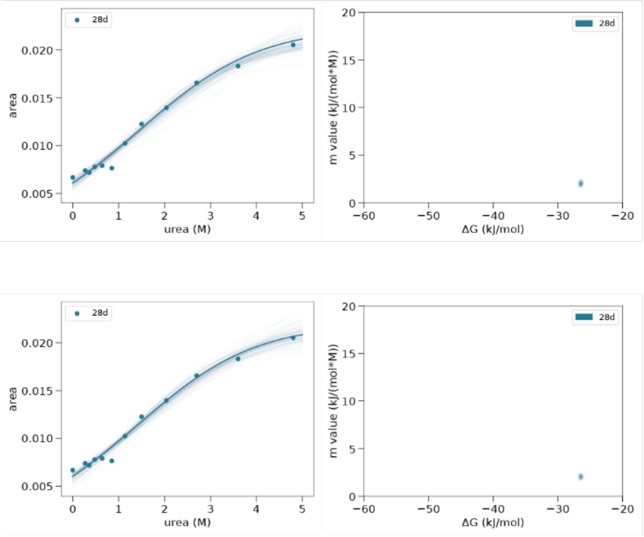


(b)

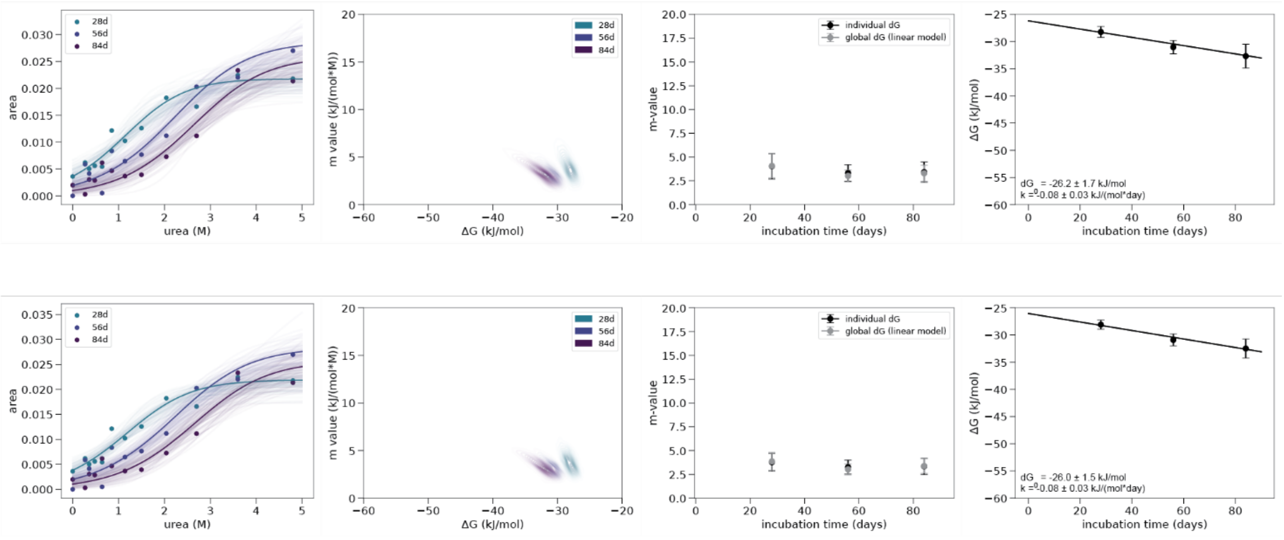


**Supplementary Figure 10.** Urea depolymerization curves of fibrils formed in condition F91 over time. (a) 1st repeat. (b) 2nd repeat. Curves were fitted using HMC sampling of solutions (n=2000) in the upper panel in (a,b) and using Bayesian analysis with a normally distributed prior on m of 3.5 with a standard deviation of 2 (lower panel in (a,b)). Column 1: Data points (area under the curve of FIDA measurements corresponding to monomeric αSyn) and their fits to the isodesmic model using HMC sampling of solutions or Bayesian analysis with HMC sampling of solutions. Column 2: Correlation of the m-value and ΔG. Column 3: Comparison of the m-value of the two different fitting approaches. Column 4: Fitted ΔG values over time (mean of 2000 solutions, error bars represent the standard deviation of the 2000 solutions) and fitting of the data to a linear model with decreasing ΔG over time: $\Delta G(t) = \Delta G_0 + k*t$. The urea depolymerization series was measured in 50 mM HEPES, pH 7.5 50 mM KCl, 5 mM $MgCl_2$, 2 mM DTT.

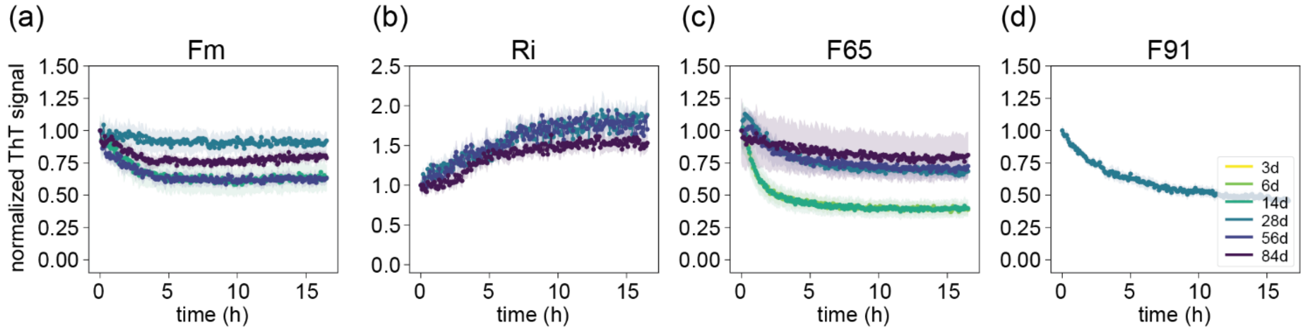


**Supplementary Figure 11.** Disaggregation of αSyn fibrils formed in condition (a) Fm, (b) Ri, (c) F65, (d) F91 over time by the tri-chaperone system HSP70, DNAJB1 and Apg2 in the presence of ATP. Fibrils were centrifuged at 16000 xg for 90 min, the supernatant was removed and the pellet containing amyloid fibrils was resuspended in the respective condition. Disaggregation of fibrils was performed in 50 mM HEPES, pH 7.5, 50 mM KCl, 5 mM $MgCl_2$, 2 mM DTT at a fibril concentration of 2 μM (monomer equivalent). Data was background corrected and normalized towards a control condition in which ATP is absent. Data corresponds to the second repeat in case of (a-c) and to the first repeat in case of (d).

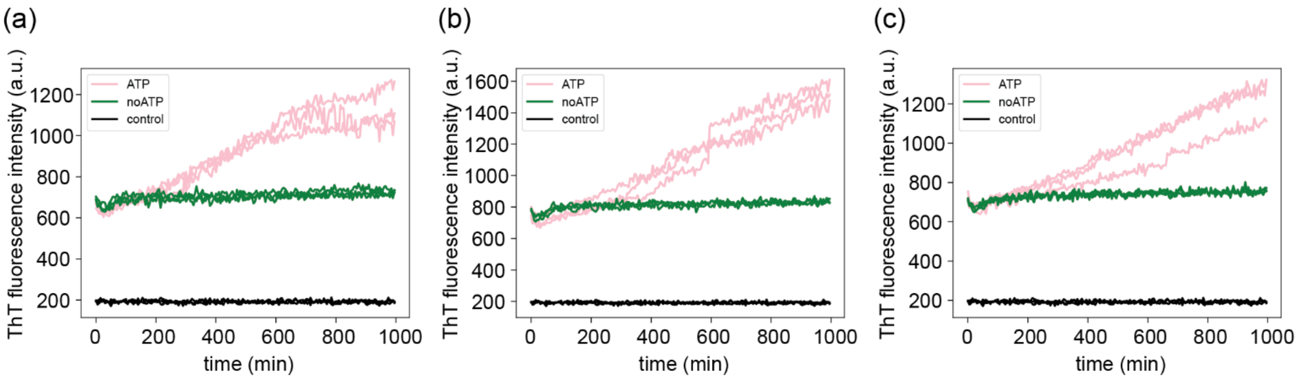


**Supplementary Figure 12.** Raw data of the disaggregation by the chaperone system Hsp70, DNAJB1 and Apg2 of fibrils formed in condition RI at (a) 28, (b) 56 and (c) 84 days of incubation of the first repeat. ThT fluorescence intensity was followed over time at 30 °C. ATP: sample containing fibrils, chaperones and ATP. noATP: sample containing fibrils and chaperones. Control: sample containing chaperones and ATP. All samples contain 20 uM ThT.

### Supporting Tables

**Supplementary Table 1.** ΔG and m-values of fitting of the isodesmic model to the depolymerization curves of the 1st repeat using Bayesian analysis with a normally distributed prior on m of 3.5 and a sigma of 2. ΔG values are in kJ/mol and m-values in kJ/(mol*M).

| day | Fm | | Ri | | F65 | | F91 | |
|---|---|---|---|---|---|---|---|---|
| | ΔG | m | ΔG | m | ΔG | m | ΔG | m |
| 3 | – | – | – | – | -26.8 ± 0.6 | 9.4 ± 1.2 | – | – |
| 6 | -28.6 ± 0.8 | 6.2 ± 1.2 | – | – | -31.1 ± 0.8 | 6.5 ± 1.0 | – | – |
| 14 | -28.4 ± 0.8 | 8.9 ± 1.4 | – | – | -30.9 ± 0.8 | 8.9 ± 1.2 | – | – |
| 28 | -34.6 ± 1.6 | 4.7 ± 0.9 | -31.3 ± 1.0 | 3.1 ± 0.7 | -30.8 ± 0.8 | 6.6 ± 0.8 | -26.5 ± 0.2 | 2.1 ± 0.3 |
| 56 | -31.4 ± 1.0 | 4.3 ± 0.7 | -36.3 ± 2.7 | 4.2 ± 1.2 | -34.0 ±1.3 | 6.7 ± 1.0 | – | – |
| 84 | -34.2 ± 2.2 | 5.2 ± 1.1 | -35.7 ± 2.3 | 2.7 ± 1.0 | -36.2 ± 2.1 | 6.3 ± 1.3 | – | – |

**Supplementary Table 2.** ΔG and m-values of fitting of the isodesmic model to the depolymerization curves of the 2nd repeat using Bayesian analysis with a normally distributed prior on m of 3.5 and a sigma of 2. ΔG values are in kJ/mol and m-values in kJ/(mol*M).

| day | Fm | | Ri | | F65 | | F91 | |
|---|---|---|---|---|---|---|---|---|
| | ΔG | m | ΔG | m | ΔG | m | ΔG | m |
| 3 | – | – | – | – | – | – | – | – |
| 6 | – | – | – | – | -31.3 ± 0.6 | 4.9 ± 0.6 | – | – |
| 14 | -36.1 ± 1.4 | 5.8 ± 0.7 | -40.0 ± 1.3 | 7.7 ± 0.7 | -31.9 ± 0.9 | 5.5 ± 0.7 | – | – |
| 28 | -31.6 ± 0.7 | 3.8 ± 0.5 | -39.6 ± 1.8 | 5.8 ± 0.8 | -33.3 ± 1.0 | 6.9 ± 0.7 | -28.1 ± 0.8 | 3.8 ± 0.9 |
| 56 | -35.4 ± 1.2 | 4.8 ± 0.7 | -41.8 ± 2.4 | 4.0 ± 1.1 | -35.6 ± 1.4 | 6.6 ± 0.8 | -30.9 ± 1.1 | 3.3 ± 0.7 |
| 84 | – | – | -40.7 ± 1.9 | 4.6 ± 0.7 | -39.9 ± 1.9 | 8.6 ± 1.0 | -32.5 ± 1.7 | 3.3 ± 0.9 |

**Supplementary Table 3.** $\Delta G_0$, k and m-values of fitting of a linear aging model $\Delta G(t) = \Delta G_0 + k*t$ to the depolymerization curves using Bayesian analysis with a normally distributed prior on m of 3.5 and a sigma of 2. m-values are in kJ/(mol*M).

| parameter | Fm | | Ri | | F65 | | F91 | |
|---|---|---|---|---|---|---|---|---|
| | 1st repeat | 2nd repeat | 1st repeat | 2nd repeat | 1st repeat | 2nd repeat | 1st repeat | 2nd repeat |
| $dG_0$ (kJ/mol) | -28.4 ± 0.75 | -32.9 ± 1.0 | -28.9 ± 1.9 | -38.9 ±1.1 | -28.6 ± 0.5 | -30.5 ± 0.6 | – | -26.0 ±1.5 |
| k (kJ/(mol*day)) | -0.08 ± 0.03 | -0.03 ± 0.03 | -0.09 ± 0.04 | -0.03 ± 0.02 | -0.09 ± 0.02 | -0.10 ± 0.02 | – | -0.08 ±0.03 |
| m-value day 3 | – | – | – | – | 12.2 ± 1.3 | – | – | – |
| m-value day 6 | 6.3 ± 1.2 | – | – | – | 4.7 ± 0.5 | 4.8 ± 0.5 | – | – |
| m-value day 14 | 10.0 ± 1.2 | 4.4 ± 0.5 | – | 7.4 ± 0.5 | 7.6 ± 0.8 | 5.5 ± 0.4 | – | – |
| m-value day 28 | 2.9 ± 0.4 | 5.0 ± 0.6 | 3.1 ± 0.7 | 5.9 ± 0.4 | 6.8 ± 0.6 | 6.9 ± 0.5 | – | 3.8 ± 0.9 |
| m-value day 56 | 5.1 ± 0.9 | 4.5 ± 0.6 | 3.1 ± 0.7 | 3.6 ± 0.6 | 6.4 ± 0.8 | 6.9 ± 0.6 | – | 3.0 ± 0.6 |
| m-value day 84 | 5.5 ± 0.9 | – | 2.6 ± 1.0 | 4.9 ± 0.5 | 6.2 ± 1.0 | 8.0 ± 0.8 | – | 3.4 ± 0.8 |

**Supplementary Table 4.** Degree of disaggregation of amyloid fibrils formed in different conditions at different time points. A disaggregation degree of one corresponds to a total loss of ThT signal while a disaggregation degree of 0 corresponds to no change in ThT intensity.

| Incubation days | Fm | | F65 | | F91 | |
|---|---|---|---|---|---|---|
| | 1st repeat | 2nd repeat | 1st repeat | 2nd repeat | 1st repeat | 2nd repeat |
| 3 | – | – | 0.93 ± 0.01 | – | – | – |
| 6 | – | – | 0.69 ± 0.01 | 0.60 ± 0.05 | – | – |
| 14 | 0.80 ± 0.13 | 0.33 ± 0.11 | 0.72 ± 0.02 | 0.59 ± 0.07 | – | – |
| 28 | 0.00 ± 0.13 | 0.06 ± 0.08 | 0.88 ± 0.01 | 0.31 ± 0.04 | 0.54 ± 0.02 | 0.55 ± 0.05 |
| 56 | 0.17 ± 0.18 | 0.34 ± 0.04 | 0.54 ± 0.04 | 0.29 ± 0.08 | – | 0.47 ± 0.03 |
| 84 | 0.20 ± 0.05 | 0.16 ± 0.04 | 0.58 ± 0.03 | 0.22 ± 0.17 | – | 0.38 ± 0.06 |
